# Regulatory Changes in Pterin and Carotenoid Genes Underlie Balanced Color Polymorphisms in the Wall Lizard

**DOI:** 10.1101/481895

**Authors:** Pedro Andrade, Catarina Pinho, Guillem Pérez i de Lanuza, Sandra Afonso, Jindřich Brejcha, Carl-Johan Rubin, Ola Wallerman, Paulo Pereira, Stephen J. Sabatino, Adriana Bellati, Daniele Pellitteri-Rosa, Zuzana Bosakova, Miguel A. Carretero, Nathalie Feiner, Petr Marsik, Francisco Paupério, Daniele Salvi, Lucile Soler, Geoffrey M. While, Tobias Uller, Enrique Font, Leif Andersson, Miguel Carneiro

## Abstract

Reptiles use pterin and carotenoid pigments to produce yellow, orange, and red colors. These conspicuous colors serve a diversity of signaling functions, but their molecular basis remains unresolved. Here, we show that the genomes of sympatric color morphs of the European common wall lizard, which differ in orange and yellow pigmentation and in their ecology and behavior, are virtually undifferentiated. Genetic differences are restricted to two small regulatory regions, near genes associated with pterin (*SPR*) and carotenoid metabolism (*BCO2*), demonstrating that a core gene in the housekeeping pathway of pterin biosynthesis has been co-opted for bright coloration in reptiles and indicating that these loci exert pleiotropic effects on other aspects of physiology. Pigmentation differences are explained by extremely divergent alleles and haplotype analysis revealed abundant trans-specific allele sharing with other lacertids exhibiting color polymorphisms. The evolution of these conspicuous color ornaments is the result of ancient genetic variation and cross-species hybridization.

## INTRODUCTION

Color morphs can be found in thousands of species in nature. In many animals, color morphs persist over extended periods of time, co-occur across large geographic regions, and differ in a range of key morphological, physiological, or behavioral traits (1-5). These features offer an exceptional framework to investigate central mechanisms of phenotypic diversity and innovation, with potential insights into a variety of processes, including adaptation, convergent evolution, sexual selection, and the early stages of sympatric speciation (1,6). It is thus of special interest to understand the evolutionary and mutational mechanisms promoting the emergence and long-term persistence of sympatric color polymorphisms in nature.

Reptiles are amongst the most colorful animals. They achieve this in different ways, but their vibrant colors spanning the gradient of hues between yellow and red are generated primarily by carotenoid and/or pterin pigments (7). The colors produced by these pigments serve many biological functions and are thought to be of key importance in intra- and interspecific communication (8). Striking colors often differ between closely related species and have been gained and lost across the reptilian phylogeny, demonstrating that coloration driven by these types of pigments can change over short time scales and presumably evolved multiple times independently. Yet, despite intense study of the biochemical basis and ecology of carotenoid- and pterin-based pigmentation in reptiles (3,7,9-11), we have a limited understanding regarding the genetic changes and molecular pathways that govern differences among individuals and species.

To investigate the genetic and evolutionary bases of the vivid colors displayed by reptiles, and to test hypothesis about how and why color polymorphisms and correlated trait variation persist within populations, we studied the European common wall lizard (*Podarcis muralis*) (Fig. 1A) – a polymorphic lizard in which the ventral scales of males and females exhibits one of three distinct colors (orange, yellow, and white) or a mosaic pattern combining two colors (orange-yellow and orange-white) (12,13). Each of these five color morphs can be found throughout most of the broad geographic distribution of the species (Fig. 1B), and are shared by intraspecific sub-lineages thought to have diverged up to 2.5 million years ago (14). While the white morph is typically the most common (>50%), the relative frequency of morphs is highly variable even at small regional scales and the yellow or orange morphs may occasionally prevail (15,16) (SI Appendix, Fig. S1). The widespread distribution and persistence of color variation is thought to be due to balancing selection and the product of an interplay between natural and sexual selection (17). Previous work has shown that morphs mate assortatively with respect to ventral color (∼75% of pairs) and differ in additional traits, including morphology, behavior, physiology, immunology, and reproduction (12,18-22). The mode of inheritance of the color morphs is unknown.

**Figure 1.**
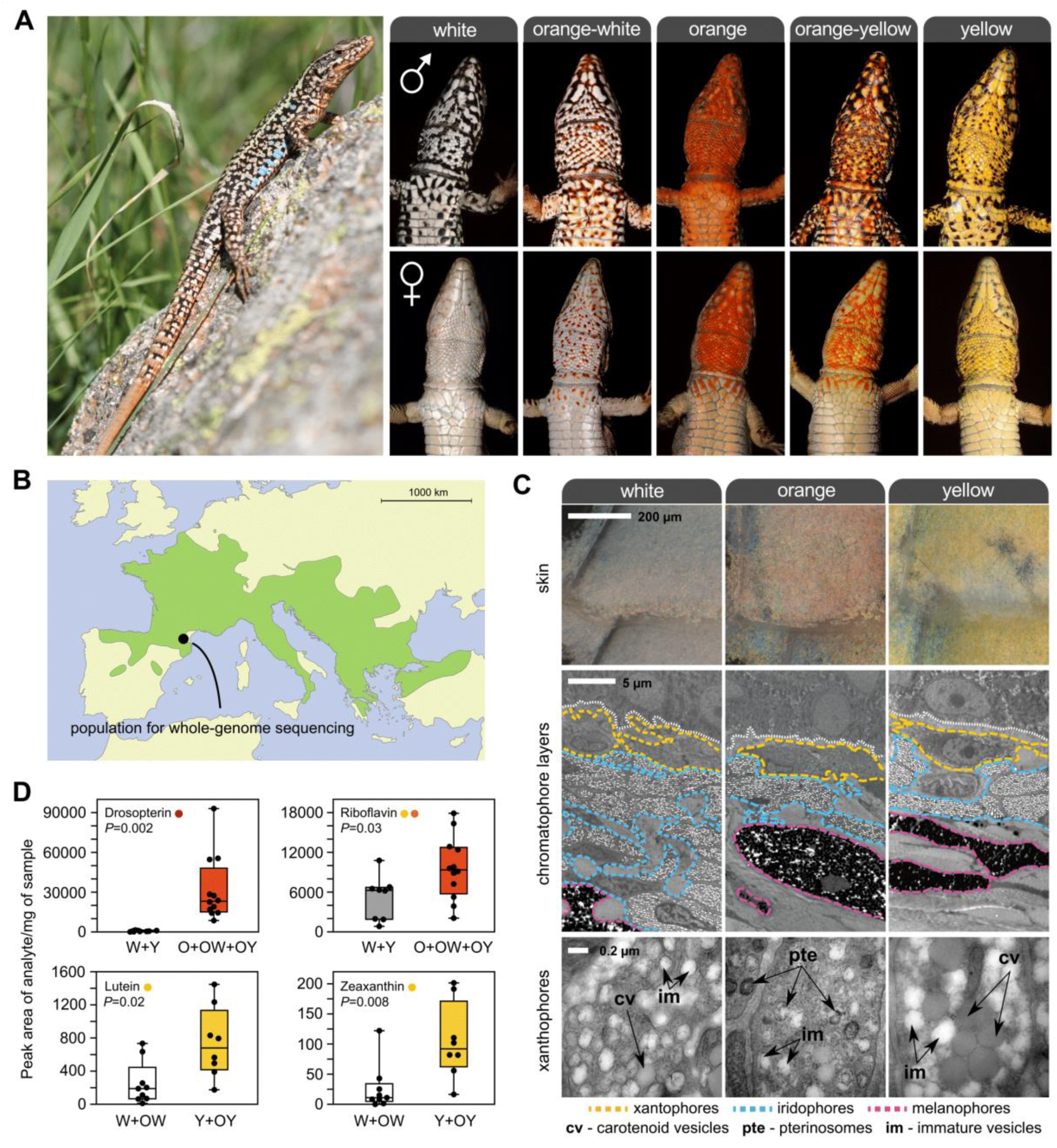
Color polymorphism in the European common wall lizard, *Podarcis muralis*. **(A)** The common wall lizard (picture on the left). The pictures on the right illustrate the five discrete ventral morphs. These conspicuous colors likely function as visual ornaments implicated in sexual signaling. The yellow and orange colors are restricted to the ventral surface and males and females exhibit marked differences in the extent of pigmentation in some populations. **(B)** Geographic distribution of the species in Europe (light green). **(C)** Ultrastructure of the ventral skin of the three pure morphs. Top panel shows a close-up of the ventral scales of each morph under a light microscope. The panel in the middle shows transmission electron microscopy of the three chromatophore layers [xantophores (yellow), iridophores (blue), and melanophores (pink)]. The bottom panel shows electromicrographs detailing the structure of xanthophores. Examples of pterinosomes (pte), carotenoid vesicles (cv), and immature vesicles (im) are highlighted. **(D)** Levels of colored pterin and carotenoid compounds in the ventral skin of the different morphs obtained by HPLC-MS/MS. W=white; O=orange; Y=yellow.

## RESULTS

### Carotenoid and pterin pigments underlie pigmentation differences

We began by determining the biochemical and cellular basis of pigmentation differences among morphs. Using electron microscopy (TEM), we found that the ventral skin of all morphs contained the same set of dermal pigment cells arranged as three superimposed layers (xantophores, iridophores, and melanophores; Fig. 1C). The iridophore layer was thinner in orange individuals compared to yellow and white, but the most noticeable difference among morphs was observed in the xantophore layer, which in reptiles usually contains pterins and carotenoid pigments (7). Although two types of morphologically distinct pigment-containing organelles (pterinosomes and carotenoid vesicles) coexisted within many xantophores irrespective of skin color, their relative abundance varied drastically. Pterinosomes were abundant in xantophores from orange skin, whereas carotenoid vesicles of large size were abundant in yellow skin. The xantophores of individuals exhibiting white skin were characterized by low numbers of pterinosomes and carotenoid vesicles, as well as by large numbers of immature vesicles of small size and low electron density under TEM.

Next, we used high-performance liquid chromatography-tandem mass spectrometry (HPLC-MS/MS) to quantify individual carotenoid and pterin derivatives (Figs. 1D, SI Appendix, Fig. S2A). In agreement with microscopy, carotenoids and pterins were present in skin of all colors, but the relative proportions of specific metabolites varied according to color morph. Both lutein and zeathanxin, two yellow-colored carotenoids, were significantly more abundant in yellow skin relative to white skin (Fig. 1D; Mann-Whitney U test, *P*<0.05 for both tests). By quantifying nine colored and colorless pterin metabolites, we found no evidence for substantial changes in the overall pterin profiles between white and yellow individuals (Figs. S2A, S2B). However, the skin of individuals expressing orange pigmentation (pure and mosaic) contained significantly higher levels of orange/red pterins (riboflavin and drosopterin) when compared to non-orange individuals (Fig. 1D; Mann-Whitney U test, *P*<0.05 for both tests). The observed variation in colored pterins and carotenoids among morphs provides a biochemical basis for the inter-individual differences in orange and yellow coloration, respectively. Together, the microscopy and biochemical analyses support the hypothesis that chromatic variation is due to alterations in the metabolism, transport, or deposition of carotenoids and pterins, or some combination of these processes.

### A highly contiguous genome sequence for the common wall lizard

As a backbone for our genetic and evolutionary studies, we sequenced and assembled a reference genome for the common wall lizard (PodMur1.0). The assembly was primarily based on Pacific Biosciences (PacBio) long-read sequences (∼100X coverage) and complemented with Illumina sequencing (30X). To aid in the assembly of the genome, we generated chromosome conformation capture sequencing data using CHiCAGO and HiC libraries (23,24). The combination of these methodologies resulted in a high-quality, chromosome-scale, genome assembly of 1.51 Gb with a contig N50 of 714.6 kb and a scaffold N50 of 92.4 Mb (SI Appendix, Table S1). Our assembly produced 19 scaffolds of large size (18 larger than 40 Mb and one larger than 10 Mb; SI Appendix, Table S2), which matches well with the karyotype of the species (2n=38) (25). A search for conserved single copy orthologues revealed that the assembly included a large percentage of full-length genes (93.2%; SI Appendix, Table S1), indicating that the sequence is both highly contiguous and accurate.

Karyotypic analysis demonstrated that the common wall lizard possesses a ZW/ZZ sex determination system and that the two sex-chromosomes are of similar size and shape (26). To identify the Z chromosome in our assembly, we compared sequence coverage between DNA pools of males and females (SI Appendix, Fig. S3). We found a single chromosome, that we named Z, for which sequence coverage in females was roughly half that of males throughout most of the chromosome, as expected from reads mapping on the Z, but not on the W chromosome. The results show that Z and W, although similar in size, have likely evolved independently for a long time and have diverged extensively in sequence. Finally, the genome assembly was annotated using *in silico* predictions and transcriptome data from five tissues and one embryonic stage (SI Appendix, Table S3), which predicted 24,656 protein-coding genes (SI Appendix, Table S1). This high-quality lizard genome provides an important resource for comparative genomic analysis within squamate reptiles.

### The distinct morphs have near-identical genome sequences

To obtain genome-wide polymorphism data, we sampled 154 individuals from two neighboring localities in the eastern Pyrenees and performed whole-genome sequencing (Fig. 1B). We sequenced 10 DNA pools of individuals grouped by color morph and locality (mean coverage=∼16X; SI Appendix, Table S4). The short sequence reads were aligned to the reference genome for variant identification and allele frequency estimation, yielding a total of 12,066,526 variants (SNPs and indels). A neighbor-joining tree based on this genome-wide data revealed that the overall genetic structure was predominantly influenced by sampling site as opposed to color phenotype (Fig. 2A). The low differentiation between morphs is well demonstrated by an average *F*_*ST*_ value close to 0 (0.02) and by a paucity of variants displaying high allele frequency differences in pairwise comparisons (SI Appendix, Fig. S4). The different morphs also displayed similar levels (*π*) and patterns (Tajima’s *D*) of nucleotide variation, as expected from individuals drawn from a single population (Fig. 2B). These results demonstrate that the genome-wide impacts of assortative mating are minor and that rates of gene flow between morphs are sufficiently high to prevent the build-up of strong genetic differentiation.

**Figure 2.**
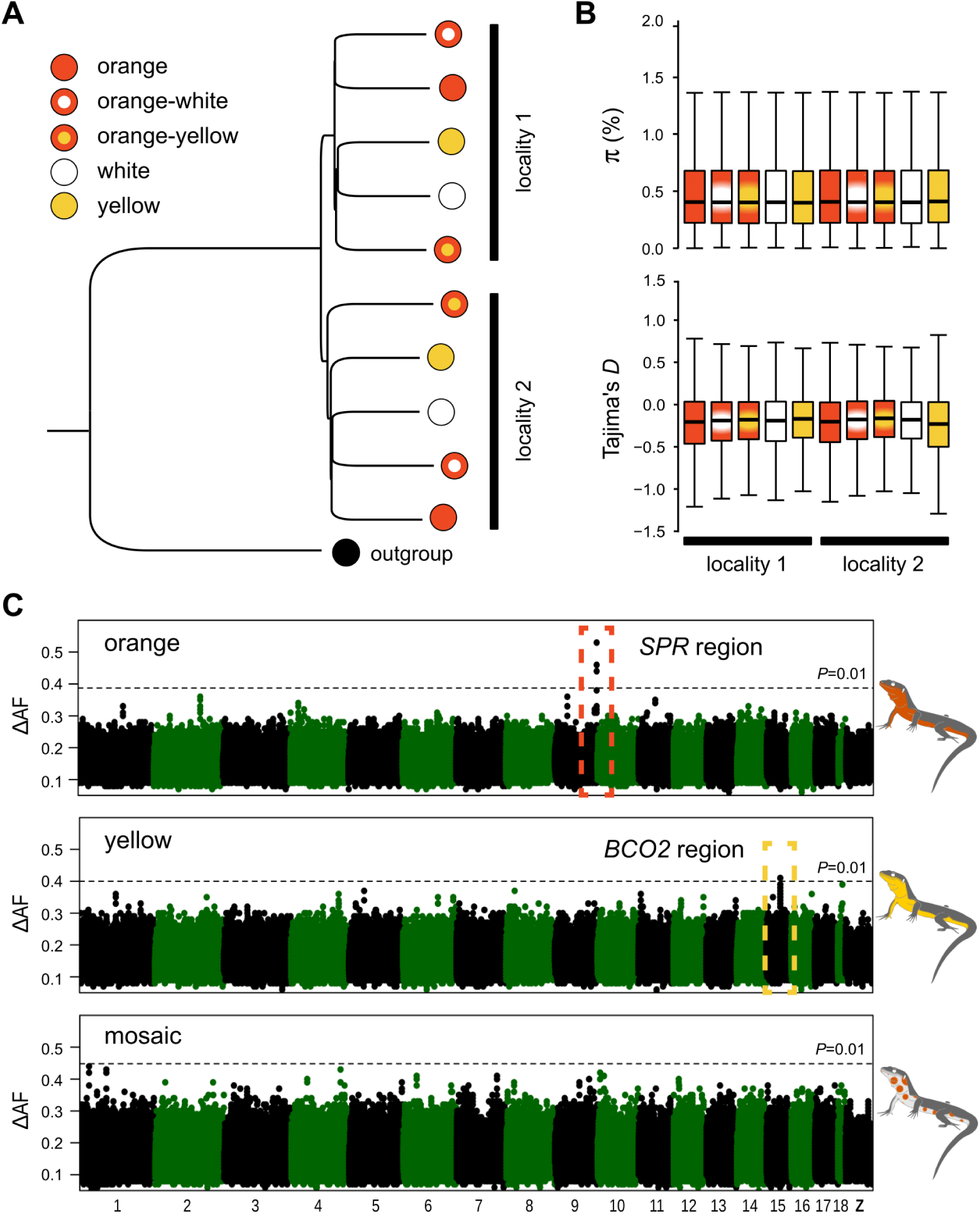
Population structure and genetic basis of color polymorphism in the common wall lizard. **(A)** Neighbor-Joining tree summarizing genetic distance among individual pools using 250,000 randomly chosen SNPs. The tree was rooted with a DNA pool of individuals sampled from Italy and belonging to a different intraspecific sub-lineage. **(B)** Nucleotide diversity (π) and Tajima’s *D* estimated for each morph. Both statistics were calculated in 10 kb non-overlapping windows and a genome-wide estimate was obtained by averaging all windows across the genome. **(C)** Genetic mapping based on differences in allele frequencies (ΔAF) for the orange, yellow, and mosaic phenotypes. The Manhattan plots show the median value of 20-SNP windows (5-SNP overlap) across the reference genome. The dashed line represents a 1% significance cut-off based on 1,000 permutations conducted for each dataset.

### High-resolution mapping of genomic regions underlying differences in coloration

Encouraged by the overall low levels of genetic differentiation among morphs, we next carried out population genomic analysis to identify chromosome regions associated with color variation. We calculated allele frequency differentiation (ΔAF) averaged in sliding windows and, guided by our phenotypic characterization, we contrasted morphs by the presence/absence of specific colors (orange and yellow) or by their patterning (mosaic). For both orange and yellow coloration, we found in each case a single genomic region showing significantly high ΔAF when compared to an empirical null distribution generated by permutation (Fig. 2C). These regions were small, located on the autosomes, and embedded within an otherwise undifferentiated genome (Fig. 3A). A Cochran-Mantel-Hanzel test (CMH), configured to identify consistent changes in allele frequency between different samples, corroborated the ΔAF analysis and revealed the same two regions as the top outliers for orange and yellow coloration (SI Appendix, Fig. S5).

**Figure 3.**
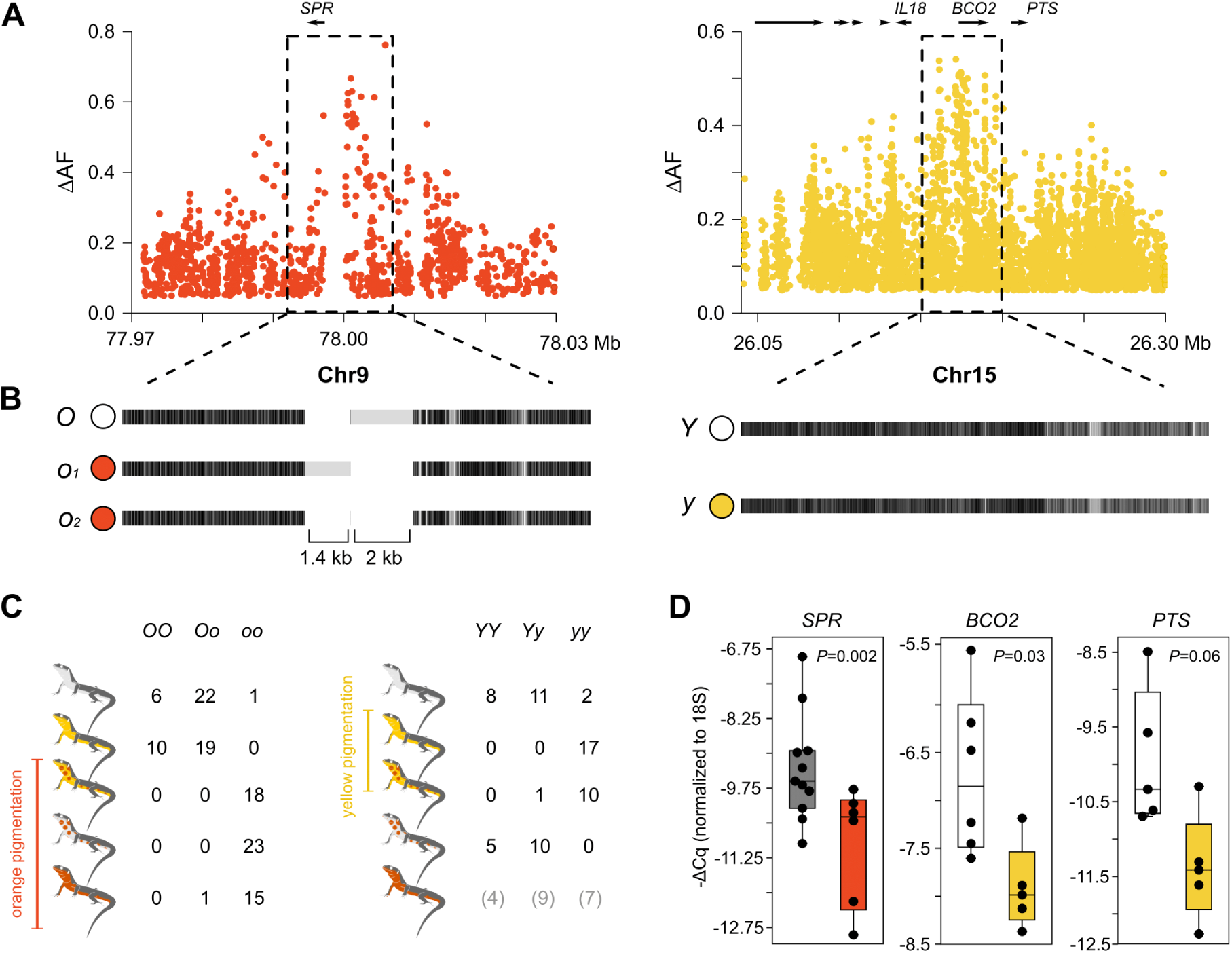
Regulatory variation explains color polymorphism in the common wall lizard. **(A)** Differences in allele frequencies (ΔAF) for the orange and yellow phenotypes around *SPR* and *BCO2* (each dot represents a SNP). **(B)** Haplotype structure for the same two regions based on the alignment of our reference genome sequence to consensus sequences of the alternative haplotypes obtained using Nanopore and Sanger sequencing. Black indicates homology and light gray indicates mismatches that can originate from point mutations or indel variants. **(C)** Individual genotypes for the *SPR* and *BCO2* loci among the five morphs based on high ΔAF variants selected from the whole-genome data. **(D)** qPCR measurements of *SPR, BCO2*, and *PTS* expression in ventral skin.

The region significantly associated with orange coloration was located near the *SPR* gene (Chr9; Fig. 3A), which encodes sepiapterin reductase, a key enzyme in pterin metabolism (SI Appendix, Fig. S6A). The presence/absence of yellow coloration mapped to the genomic region around the *BCO2* gene (Chr15; Fig. 3A). *BCO2* encodes the beta-carotene oxygenase 2 that oxidizes colorful carotenoids to colorless apocarotenoids during the biosynthesis of vitamin A (SI Appendix, Fig. S6B). In light of the phenotypic differences between morphs, *PTS* is another gene of interest that was located near the *BCO2* region (Fig. 3A). *PTS* encodes the 6-pyruvoyltetrahydropterin synthase, which is also a key enzyme in pterin metabolism (SI Appendix, Fig. S6A). Although evolutionary theory predicts that alleles encoding sexually selected traits such as color ornaments should accumulate preferentially on sex-chromosomes (27-28), particularly for ZZ/ZW systems, our results demonstrate that the two genomic regions most strongly associated with orange and yellow pigmentation are not sex-linked.

To confirm the association and gain additional insight about the genetic architecture of orange and yellow coloration, we obtained individual genotypes for high ΔAF variants selected from the resequencing data (Fig. 3C). This genotyping revealed that, in 56 out of 57 cases, orange and mosaic individuals were homozygous for one allele (hereafter *o*) at the *SPR* locus, whereas yellow and white individuals, with one exception, were either heterozygous (n=41) or homozygous (n=16) for the alternative allele (hereafter *O*) – a highly significant association (recessive model, *P=*9.3×10^−29^). The association at the BCO2 locus was also highly significant (recessive model, *P=*1.2×10^−15^) with most individuals displaying yellow coloration (yellow and orange-yellow) being homozygous (27 out of 28) for one allele (hereafter *y*), and 34 out of 36 individuals displaying white coloration (white and orange-white) being heterozygous (n=21) or homozygous (n=13) for the alternative allele (hereafter *Y*). The few discordant individuals in both loci can be explained by incomplete linkage between the genotyped and the causal variants, incomplete penetrance due to environmental effects, or by other interacting genetic factors. We found the three different genotypic classes among individuals of the pure orange morph at the *BCO2* locus, suggesting that yellow carotenoids, if present, are likely masked by the stronger pterin-based orange coloration. While other loci not revealed in our genomic analysis might play a minor role, these loci explain a large component of the differences in pterin and carotenoid pigmentation among morphs.

Subsequently, we extended the genotyping to a larger cohort of common wall lizards covering the broad geographic range of the species and belonging to several of its intraspecific sub-lineages (14). This analysis confirmed that *SPR* is the primary locus explaining orange coloration across the species’ range (SI Appendix, Table S5). In contrast, the variant near *BCO2*, which was strongly associated with yellow pigmentation in the Pyrenees, was either not present in other sub-lineages or not evidently associated with yellow coloration (SI Appendix, Table S5). The absence of an association might be explained by some of the same reasons as above, or by the possibility that yellow coloration might have evolved convergently more than once, either through independent mutation events near *BCO2* or through a different gene.

By contrast, our genomic analysis (ΔAF and CMH) did not reveal any genetic candidate associated with mosaic morphs (Fig. 2C, SI Appendix, Fig. S5). The mosaic morphs are also not explained by any combination of alleles between *SPR* and *BCO2* (Fig. 3C). The lack of genetic signal could be explained by a polygenic architecture consisting of a few loci with small-moderate effects that our sequencing efforts might not be powerful enough to detect. Alternatively, this phenotype might not be directly under genetic control, but rather reflect a temporary ontogenetic stage or phenotypic plasticity. In fact, a long-term study of a natural population showed that many sub-adult individuals displaying isolated orange scales, identical to the orange-white morph, subsequently develop pure orange coloration (12). Additional work is needed to resolve the genetic basis (or lack thereof) of the mosaic morphs.

### Synteny and gene content are preserved between haplotypes associated with coloration

Structural changes could be responsible for the observed genetic differentiation associated with each color morph. To explore this possibility, we sequenced an additional individual homozygous for the haplotypes not represented in the reference sequence (*o* and *Y*) using Nanopore long reads (9.7X coverage). We found that the region immediately upstream of *SPR* (∼7 kb) contained two medium-size indel polymorphisms when compared to the reference sequence (1.4 and 2 kb) (Fig. 3B). A PCR amplification of this region in a larger number of orange individuals revealed two orange alleles of different size (*o1* and *o2*) that shared the 2 kb indel (Fig. 3B). However, we failed to identify homologous sequences to the two indels in the chicken or the anole lizard, thus they are likely derived insertions lacking any strongly conserved sequence. In the region associated with yellow coloration surrounding *BCO2*, the *y* and *Y* haplotypes were identical in their general structure (Fig. 3B). Taken together, we found no evidence for the existence of large-scale structural differences, copy number variants, or translocations in both loci. Thus, gene content is preserved between the haplotypes that control pigmentation differences, demonstrating that the described multi-trait divergence between morphs is not explained by a supergene organization implicating many genes.

### Cis-regulatory sequence variation underlies the color polymorphism

Next, we examined in greater detail the region of association around *SPR* and *BCO2*. The signal of association for orange coloration was restricted to a noncoding region immediately upstream of *SPR* (Fig. 3A). The association for yellow coloration in *BCO2* was strong both upstream and overlapping part of the coding region (Fig. 3A); however, we found no mutations strongly associated with yellow coloration that could alter the protein structure of *BCO2* (nonsynonymous, stop, splicing, or frameshift). We thus hypothesized that the pigmentation differences are controlled by regulatory sequence variation at these loci.

To examine the role of regulatory variation in pigmentation differences, we studied gene expression in several tissues (ventral skin, brain, muscle and liver) harvested from orange, yellow, and white lizards using quantitative PCR (qPCR) (Fig. 3D, SI Appendix, Fig. S6C). *SPR* expression was significantly lower in orange skin relative to other colors (Mann-Whitney U test, *P=*0.002), whereas the same was found for *BCO2* in yellow skin relative to white skin (Mann-Whitney U test, *P*=0.03). Low *BCO2* expression is consistent with reduced activity of this enzyme that presumably leads to accumulation of colorful carotenoids, similarly to what has been shown for carotenoid-based yellow coloration in birds (29). Despite the fact that the strongest signal of association was located near *BCO2* (Fig. 3A), the expression of *PTS* was also higher in white individuals when compared to yellow individuals, although not significantly so (Mann-Whitney U test, *P*=0.06). Given that we found no obvious differences in pterin content between white and yellow individuals (SI Appendix, Fig. S2A, 2B), it is unclear whether *PTS* could play a role in pigmentation differences. In the other tissues, we found no significant differences between morphs for *BCO2*, whereas *SPR* was slightly upregulated in muscle and brain of orange individuals – the opposite pattern that was observed in the skin – and the differences were significant for the muscle and marginally significant for the brain (SI Appendix, Fig. S6C).

We next examined the expression differences in *BCO2* and *SPR* by measuring allele-specific expression in the skin. If cis-acting regulatory mutations are driving expression differences between morphs, then one haplotype should preferentially be expressed in the shared cellular environment of individuals heterozygous for the *Y/y* or *O/o* alleles. Among the individuals used in the qPCR analysis, we found for each gene one heterozygous individual containing a polymorphism overlapping the coding region that could be used to quantify allelic imbalance using cDNA sequencing. Corroborating the observed differences in the qPCR experiments, we found a strong preferential expression of one allele over the other, both for *SPR* (81% vs. 19%; n=8,210 reads; Chi-square, *P*<10^−16^) and *BCO2* (72% vs. 28%; n=25,026 reads; Chi-square, *P*<10^−16^). Overall, our expression studies provide evidence that pigmentation differences in the skin evolved through one or more cis-acting mutations affecting the activity of *SPR* and *BCO2*, and possibly *PTS*. They further suggest that gene expression of *SPR* in other tissues might also differ between morphs.

### Genetic variation upstream of *SPR* and *BCO2* is shared among divergent species

To gain insight into the evolutionary history of the two regions that explain pterin and carotenoid-based pigmentation, we sequenced amplicons (∼550 bp) overlapping the strong signals of association upstream of *SPR* and *BCO2* in individuals from the Pyrenees, and used the sequence data to construct haplotype trees (Fig. 4A). Both in *BCO2* and *SPR*, the haplotypes typically associated with presence or absence of yellow (*y* and *Y*) and orange coloration (*o* and *O*) clustered into extremely divergent haplogroups (Fig. 4A). In *SPR*, we also uncovered two divergent haplotypes associated with non-orange coloration. The average number of pairwise differences (*d*_*XY*_) between the orange and non-orange haplogroups in *SPR* (*d*_*XY*_ = 5.2-6.9%), and between the yellow and non-yellow haplogroups in *BCO2* (*d*_*XY*_ = 3.1%), was ∼8-17-fold higher than the average number of pairwise differences between sequences within the French population (*π*≈0.4%; Fig. 2B). To put this into context, the average human-chimpanzee nucleotide divergence in non-coding regions is ∼1.2% (30). The magnitude of divergence between haplotypes associated with pigmentation differences show that the alleles controlling the presence or absence of orange and yellow coloration are evolutionary old and not the result of recent mutational events.

**Figure 4.**
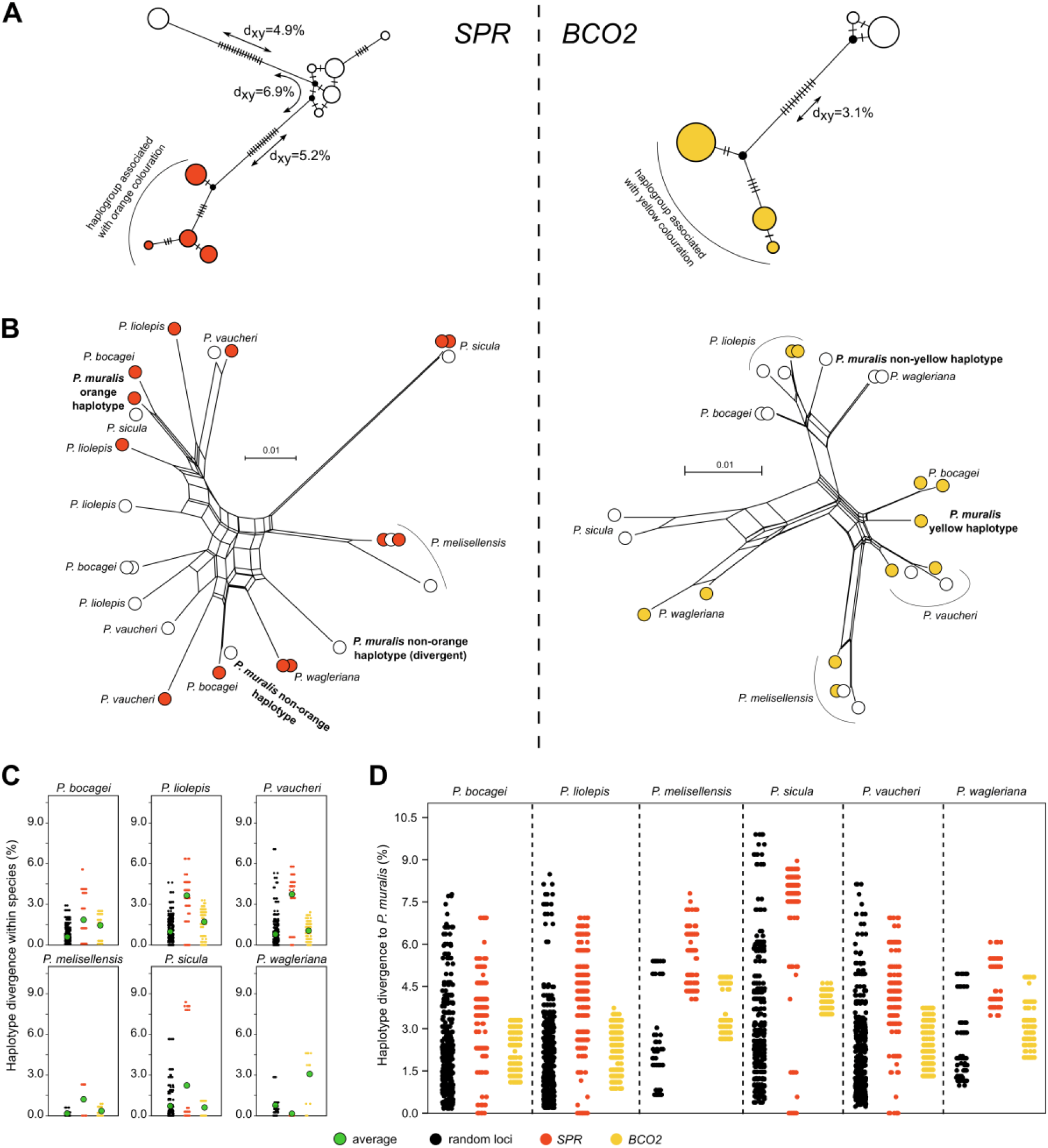
Evolution of the regulatory regions associated with pterin and carotenoid-based coloration in the genus *Podarcis*. **(A)** Median-joining genealogies of the genomic regions associated with coloration upstream of *SPR* (left) and *BCO2* (right) from common wall lizards from the Pyrenees. The dashes on branches indicates the number of observed mutations. **(B)** NeighborNet trees of the genomic regions associated with coloration upstream of *SPR* (left) and *BCO2* (right) combining common wall lizards from the Pyrenees and six other species in the genus *Podarcis*. Haplotypes are colored orange, yellow, or white, indicating the color morph of the individual. For representation purposes, only a subset of the sequences is presented. **(C)** Pairwise nucleotide differences between haplotypes within species for *SPR* (orange), *BCO2* (yellow), and 31 random loci (black). The average is indicated by a green circle. **(D)** Pairwise nucleotide differences between haplotypes of the common wall lizard and other *Podarcis* species for *SPR* (orange), *BCO2* (yellow), and 31 random loci (black). The contrasts involving other species are presented in SI Appendix, Fig. S7.

Other European lacertids exhibit intraspecific color polymorphisms resembling that of the common wall lizard (12). To study the evolutionary history of the regulatory regions associated with pigmentation in an extended phylogenetic context, we sequenced the same amplicons for six additional species of the genus *Podarcis* showing color polymorphism. Both in *SPR* and *BCO2*, we found a slight non-random clustering with respect to color morph across species. However, assuming the same recessive pattern of inheritance inferred for the common wall lizards, we would not expect to see haplotypes derived from orange and yellow individuals clustering together with haplotypes from non-orange and non-yellow individuals. The discrepancy could be explained by a combination of incomplete linkage between the sequenced amplicons and the causative mutations, as well as independent molecular bases for orange and yellow coloration in some species. Notably, the genealogies at both loci revealed abundant trans-species polymorphism, in which haplotypes from a particular species were more closely related to haplotypes from a different species (Fig. 4B). For both loci, the average number of pairwise differences between haplotypes within taxa for several *Podarcis* species was also elevated when compared to a set of 31 amplicons randomly distributed across the genome (average size = 410 bp; Fig. 4C). This pattern was particularly strong at *SPR* where a subset of the comparisons exceeds all values for the random loci, demonstrating that highly divergent haplotypes also co-occur in these species.

The patterns of deep haplotype divergence and allele sharing between species at both pigmentation loci could be explained by ancient alleles that predate speciation events and/or by interspecific hybridization. These scenarios generate different predictions regarding patterns of molecular variation. For example, introgressed haplotypes are expected to show reduced sequence divergence between species compared to the genome-wide average, whereas the ancestral polymorphism scenario predicts no such reduction. We compared pairwise haplotype differences between species for *SPR, BCO2*, and the 31 random loci (Fig. 4D; SI Appendix, Fig. S7). In contrasts between the common wall lizard and other species, the distribution of haplotype divergence for *BCO2* was within the distribution of values calculated for the random loci, consistent with a scenario of ancestral genetic variation preserved between species. In the case of *SPR*, however, we found identical haplotypes between the common wall lizard and *P*. *bocagei, P*. *liolepis, P*. *sicula*, and *P*. *vaucheri*, a pattern never observed for the random loci (Fig. 4D). The same pattern was observed in other interspecific contrasts (SI Appendix, Fig. S7). Given that these species have diverged from the common wall lizard for millions of years (>10 Mya) (31), these identical sequences strongly suggest that introgression via hybridization occurred with these or related species. Collectively, these results suggest that patterns of molecular evolution at the pigmentation loci are explained by a combination of ancestral genetic variation and introgression.

## DISCUSSION

Reptiles exhibit striking variation in color within and between species. Here, we combined whole-genome sequencing with gene expression, biochemical, and microscopy analyses to dissect the molecular basis of the coloration differences and correlated traits among morphs of the common wall lizard. Our analysis revealed, for the first-time, genes responsible for production of carotenoid- and pterin-based pigmentation in reptiles, providing the basis to unravel the evolutionary processes which influenced the emergence of the broad palette of colors and visual ornaments so prevalently found in the reptilian world. Co-option – the recruitment of genes and gene pathways to serve a different function – is emerging as a major evolutionary force underlying pigmentation novelty (32). Examples include a cytochrome P450 enzyme required for color vision and implicated in carotenoid-based red coloration in birds (33,34), a polyketide synthase essential for producing yellow psittacofulvins in parrots (35), and early-developmental genes governing wing patterns in butterflies (36). Pterins are produced through ancient multistep biosynthetic pathways and orthologs of *SPR* exist throughout the animal kingdom. The SPR enzyme is required to catalyze one of the three key steps in the synthesis of tetrahydrobiopterin (SI Appendix, Fig. S6A), an essential cofactor for a vast range of enzymatic reactions of key metabolic importance, including the degradation of the amino acid phenylalanine and biosynthesis of the neurotransmitters dopamine, serotonin, melatonin, noradrenalin and adrenalin (37). For example, defects in pterin metabolism in humans are associated with many neurological, behavioral, and movement disorders (38). Given that *SPR* has housekeeping functions and is expressed constitutively across skin and other tissues in vertebrates that do not use pterin-based compounds in pigmentation (39), our results show that a key enzyme in pterin metabolism such as SPR, and possibly PTS, has been co-opted for color variation through changes in gene regulation.

This study further expands our understanding of the genetic mechanisms linking color polymorphism to other phenotypic and fitness-related traits. It is well established that chromosomal rearrangements, such as inversions, can facilitate the evolution of multi-trait differences by suppressing recombination and preserve tight physical linkage over extended genomic segments (40-43). In contrast, genetic divergence between color morphs in the common wall lizard was restricted to very small and localized genomic intervals adjacent to pigmentation genes. This important result raises an immediate question: how do the described differences in morphology, physiology and behavior between morphs arise from such a simple genetic architecture? One plausible explanation is pleiotropic effects of the loci controlling skin pigmentation on other biological processes. The alleles associated with pigmentation differences were characterized by striking sequence divergence and it is therefore possible that each variant allele carries multiple mutations affecting the expression of *SPR, BCO2* and, possibly, *PTS*, differently in skin and other tissues, for instance in brain regions controlling behavioral differences between morphs. Carotenoid and pterin compounds are known to be involved in a wide range of vital metabolic processes (37,44), thus, alterations in the regulation of their biosynthetic pathways may form the basis of the multi-trait divergence often observed between color morphs in many species of reptiles.

Finally, we show that trans-specific allele sharing in both pigmentation loci was frequent among several species harboring identical color polymorphisms to the common wall lizard. Our results indicate that a combination of balancing selection and introgressive hybridization played a role in the evolution of coloration in the genus *Podarcis* and led to an unusually long-term maintenance of genetic variation at these pigmentation loci. In the last few years, studies have revealed that introgression can transfer adaptive traits and evolutionary novelty between populations and species (45-47). Since polymorphic ventral coloration is common in lacertid lizards from multiple genus other than *Podarcis*, our results raise the intriguing possibility that hybridization might be a frequent mechanism promoting cross-species transfer of visual ornaments and associated traits, which are subsequently kept by balancing selection within species. As genomic data accumulate, the molecular determinants of color variation in other polymorphic systems and the role of hybridization and ancestral variation in the evolution of visual ornaments can be evaluated systematically.

## METHODS

Detailed methods are available as Supporting information.

## ACKNOWLEDGMENTS

This work was supported by research grants from the Knut and Alice Wallenberg Foundations (to LA); by the Swedish Research Council (to LA [2017-02907] and TU [E0446501]); by the Fundação para a Ciência e Tecnologia (FCT) through POPH-QREN funds from the European Social Fund and Portuguese MCTES (FCT Investigator grants to MC [IF/00283/2014/CP1256/CT0012] and CP [IF/01597/2014/CP1256/CT0009], and post-doc fellowships to SS [SFRH/BPD/99138/2013] and GPL [SFRH/BPD/94582/2013]); by a Wallenberg Academy Fellowship to TU; by research fellowships attributed to PA (PD/BD/114028/2015) and PP (PD/BD/128492/2017) in the scope of the Biodiversity, Genetics, and Evolution (BIODIV) PhD program at CIBIO/InBIO and University of Porto; by the project “PTDC/BIA-EVL/30288/2017 - NORTE -01-0145-FEDER-30288”, co-funded by NORTE2020 through Portugal 2020 and FEDER Funds and by National Funds through FCT; by the projects “Genomics and Evolutionary Biology” and “Genomics Applied to Genetic Resources” co-financed by North Portugal Regional Operational Programme 2007/2013 (ON.2 – O Novo Norte) under the National Strategic Reference Framework (NSRF) and the European Regional Development Fund (ERDF); and by an EU FP7 REGPOT grant (CIBIO-New-Gen, 286431). MAC was supported by the project NORTE- 01-0145-FEDER-000007. We thank J. Ábalos, A. Badiane, and S. Reguera for assistance during field work, and Alvarina Couto, Sara Rocha, Carla Luís, Carolina Pereira, and Joana Mendes for preliminary analyses of the 31 random amplicons. Permits for capturing and sacrificing lizards were provided by the Prefecture des Pyrénées Orientales (Arrêté n° 2016- 2-09) and the Servei de Biodiversitat i Protecció dels Animals (SF/474).

## SUPPORTING INFORMATION

## METHODS

### Phenotypic characterization

#### Microscopy

The ultrastructure, distribution, and relative abundance of chromatophores was studied with light and transmission electron microscopy (TEM) following standard procedures (1). Pieces of skin (ca. 2 mm^2^) from focal regions (throat, belly, and ventral tail) of all morphs of *P*. *muralis* (orange, n=7; white, n=7; yellow, n=4; orange-yellow, n=1; orange-white, n=1), sampled from Llívia (42°27’N, 1°58’E), were excised immediately after sacrifice, placed in Karnovsky’s fixative (2% paraformaldehyde, 2.5% glutaraldehyde in 0.1 M PB buffer, pH = 7.3) and stored at 4°C. Samples were washed with 0.1M PB, postfixed with 2% osmium tetra oxide (in 0.1M PB solution), washed with 2% uranyl acetate in 70% ethanol solution, dehydrated in an increasing ethanol series, washed in propylenoxid solution, and transferred to resin (Durcupan, Sigma). Polymerized resin blocks were cut on a Leica UCT Ultracut ultramicrotome. Ultra-thin sections were put on mesh grids, stained with lead citrate, and both observed and photographed on a FEI Tecnai Spirit G2 TEM equipped with a digital camera (Soft Image System, Morada) and image capture software (ITEM). Magnification at TEM ranged from 1,250 to 43,000x depending on the structures observed. The intensity of the electron beam was adjusted to be in the optimal range for different magnifications.

#### Biochemical analyses

The carotenoid and pteridine content in the skin of all morphs was determined using chromatographic methods. Samples of integument (ca. 12 mm^2^) from the throat and belly of individuals of each morph (orange, n=4; white, n=4; yellow, n=5; orange-yellow, n=4; orange-white, n= 5) were excised, immediately cleaned mechanically to remove muscle and connective tissue, washed briefly with PB to get rid of potential contamination from blood or body fluids, divided in two halves to be analyzed separately for carotenoids and pteridine derivatives, and frozen at −20°C until analyses.

Carotenoids were extracted with 0.5 mL ethyl acetate for 4 days at room temperature in complete darkness. The extracts were then evaporated to dryness by a stream of nitrogen at 27°C and stored at −18°C. Prior to the analyses, samples were diluted in 200µl ethyl acetate. Standards of lutein, zeaxanthin, asthaxanthin, and canthaxanthin were purchased from Sigma Aldrich (Munich, Germany). Stock solutions of the external standards were prepared at a concentration of 0.1 mg/mL by dissolving in ethyl acetate. Carotenoids were determined using UPLC coupled with PDA and MS detector. The UPLC system Dionex Ultimate 3000 (Thermo Fischer, USA) consisted of autosampler, binary pump, and diode-array (PDA) detector. The column (Kinetex C18 RP, 2.6 mm, 150 × 2.1 mm; Phenomenex, USA) maintained at 35°C was used for separation. Acetonitrile (A), methanol/water 1:1 v/v (B) and a mixture of tert-Butyl methyl ether/acetonitrile/methanol 86:86:8 v/v/v (C) were used as mobile phases for gradient elution with a constant flow rate of 0.2 mL/min. Chromatograms were monitored at 445 and 472 nm. The identity of the carotenoids was confirmed by High Resolution Accurate Mass (HRAM) Q-TOF mass spectrometer (IMPACT II, Bruker Daltonik, Germany) coupled to the UPLC/PDA system. Samples were measured with electrospray (ESI), as well as atmospheric pressure chemical ionization (APCI) in positive mode.

Pteridine derivatives were extracted with dimethyl sulfoxide (DMSO) following a previously published procedure (2). Standards of 6-biopterin, D-neopterin, leucopterin, pterin, pterin-6-carboxylic acid, and riboflavin were purchased from Sigma Aldrich (Munich, Germany). Isoxanthopterin and xanthopterin were obtained from Fluka (Buchs, Switzerland). Drosopterin was prepared following (3). Stock solutions of the external standards were prepared at a concentration of 0.1 mg/mL by dissolving them in DMSO. The working solution of the mixture of all the studied pteridine derivatives in DMSO was prepared at a concentration of 0.01 mg/ml from the stock solutions. All chromatographic measurements were carried out on a HPLC system Agilent series 1290 coupled with a Triple Quad 6460 tandem mass spectrometer (Agilent Technologies, Waldbronn, Germany). For data acquisition, the Mass Hunter Workstation software was used. A ZIC^®^-HILIC (4.6 × 150 mm, μm) column, based on zwitterionic sulfobetaine groups, was chosen (Merck, Darmstadt, Germany). The chromatographic and detection conditions were adapted from (4). The isocratic elution at a flow rate of 0.5 ml/min with the mobile phase consisted of acetonitrile/5mM ammonium acetate buffer, pH 6.80 at a volume ratio of 85:15 (*v/v*), was used for the separation of all the studied compounds with exception of drosopterin. Since drosopterin exhibits higher polarity compared with other pterin compounds, we performed a run at a volume ratio 55:45 (*v/v*) and flow rate of 0.8 ml/min. The tandem mass spectrometry (MS/MS) measurements were performed in the selected reaction monitoring mode (SRM) with positive ionization. SRM conditions used for MS/MS detection are listed in SI Appendix, Table S6. For compounds marked by an asterisk, the MS/MS conditions were adopted from (4).

### Reference genome sequencing and assembly

#### Long read sequencing and assembly

To generate a reference genome sequence for the common wall lizard, we sequenced a yellow male individual from the Pyrenees region (Llívia, 42°27’N, 1°58’E) using Pacific Biosciences (PacBio) single molecule real time sequencing (SMRT). After dissection, the different tissues were stored at −80°C until DNA preparation. Pure genomic DNA (10 ug) was obtained from muscle tissue and fragmented to 20 kb using Hydroshear DNA shearing device (Digilab). The sheared fragments were size-selected for 7-50 kb size window on Blue Pippin (Sage Science). The sequencing libraries were prepared following the standard SMRT bell construction protocol and sequenced on 100 PacBio RSII SMRT cells using the P6-C4 chemistry. Raw data was imported into SMRT Analysis software 2.3.0 (PacBio) and filtered for subreads longer than 500 bp or with polymerase read quality above 75. A *de novo* assembly of filtered subreads was generated using *FALCON* assembler version 0.4.0 (5). We used the following configuration file:

*input_fofn = input*.*fofn input_type = raw length_cutoff = 8000*

*length_cutoff_pr = 8000*

*pa_concurrent_jobs = 24*

*cns_concurrent_jobs = 36*

*ovlp_concurrent_jobs = 36*

*pa_HPCdaligner_option = -v -dal64 -t16 -e*.*70 -l1000 -s1000 ovlp_HPCdaligner_option = -v -dal64 -t32 -h60 -e*.*96 -l500 -s1000 pa_DBsplit_option = -x500 -s200*

*ovlp_DBsplit_option = -x500 -s200*

*falcon_sense_option = --output_multi --min_idt 0*.*70 --min_cov 4 --local_match_count_threshold 2 -*

*-max_n_read 200 --n_core 8 --output_dformat*

*overlap_filtering_setting = --max_diff 100 --max_cov 100 --min_cov 1 --bestn 10 --n_core 8*

#### Short read sequencing and assembly correction

To improve the accuracy of the genome sequence generated by *FALCON*, we sequenced the same individual at 30X coverage using Illumina reads. The assembly was corrected using *Pilon* (6) after mapping the short Illumina reads against the contigs obtained from the PacBio sequencing by means of *bwa-mem* (v0.7.5a-r405) (7).

#### Chicago library preparation and sequencing

Two Chicago libraries from an additional individual sampled from the same locality were prepared by Dovetail Genomics (https://dovetailgenomics.com/) as described by (8). Unlike HiC libraries (see below), Chicago libraries are generated from proximity ligation of chromatin assembled in vitro rather than chromatin obtained from in vivo sources. This reduces confounding biological signal, such as telomeric clustering or chromatin looping. Briefly, for each library, ∼500 ng of high molecular weight genomic DNA (mean fragment length = 150 kb) was reconstituted into chromatin *in vitro* and fixed with formaldehyde. Fixed chromatin was digested with *DpnII*, the 5’ overhangs filled in with biotinylated nucleotides, and then free blunt ends were ligated. After ligation, crosslinks were reversed and the DNA purified from protein. Purified DNA was treated to remove biotin that was not internal to ligated fragments. The DNA was then sheared to ∼350 bp mean fragment size and sequencing libraries were generated using NEBNext Ultra enzymes and Illumina-compatible adapters. Biotin-containing fragments were isolated using streptavidin beads before PCR enrichment of each library. The libraries were sequenced on an Illumina platform. The number and length of read pairs produced for each library was: 122 million, 2×151 bp for library 1; 145 million, 2×151 bp for library 2. When combined, these Chicago library reads provided 437.8X physical coverage of the genome (1-50 kb pairs).

#### HiC library preparation and sequencing

Two HiC libraries from the same individual used for the Chicago libraries were prepared by Dovetail Genomics as described by (9). Briefly, chromatin was fixed in place with formaldehyde in the nucleus and then extracted, for each library, independently. Fixed chromatin was digested with *DpnII*, the 5’ overhangs filled in with biotinylated nucleotides, and then free blunt ends were ligated. After ligation, crosslinks were reversed and the DNA purified from protein. Purified DNA was treated to remove biotin that was not internal to ligated fragments. The DNA was then sheared to ∼350 bp mean fragment size and sequencing libraries were generated using NEBNext Ultra enzymes and Illumina-compatible adapters. Biotin-containing fragments were isolated using streptavidin beads before PCR enrichment of each library. The libraries were sequenced on an Illumina HiSeq platform. The number and length of read pairs produced for each library was: 104 million, 2×151 bp for library 1; 134 million, 2×151 bp for library 2. These HiC library reads provided 11,977.9X physical coverage of the genome (1-50kb pairs).

#### Scaffolding the assembly with HiRise

The input *de novo* assembly, shotgun reads, Chicago library reads, and HiC library reads, were used as input data for *HiRise* software, a software pipeline designed specifically for using proximity ligation data to scaffold genome assemblies (8). This analysis was carried out by Dovetail Genomics. An iterative analysis was conducted. First, Shotgun and Chicago library sequences were aligned to the draft input assembly using a modified *SNAP* read mapper (http://snap.cs.berkeley.edu). The separations of Chicago read pairs mapped within draft scaffolds were analyzed by *HiRise* to produce a likelihood model for genomic distance between read pairs, and the model was used to identify and break putative misjoins, to score prospective joins, and make joins above a threshold. After aligning and scaffolding Chicago data, Dovetail HiC library sequences were aligned and scaffolded following the same method. After scaffolding, PacBio shotgun sequences were used to close gaps between contigs. Finally, in order to improve the sequence accuracy of the gapped filled assembly, two rounds of sequence polishing were performed using the *Arrow* consensus calling algorithm (https://github.com/PacificBiosciences/GenomicConsensus). Scaffolds were named as chromosomes from larger to smaller, except for the Z-chromosome (see below).

#### Sex-chromosome identification

Lacertid lizards such as *P*. *muralis* are known to possess genetic sex determination (ZZ/ZW) (10). To identify the Z-chromosome, we compared sequence coverage between males and females using DNA pools. We sampled 24 individual females (with a balanced representation of the different morphs) from one of the localities used for the mapping of the color morphs (locality 2, SI Appendix, Table S4 and see section “Whole-genome resequencing of color morphs” below for details on DNA and library preparation). The resulting library was sequenced to an average coverage of 65.5X (SI Appendix, Table S4). To create a pool of male samples we merged all sequence alignment files generated from locality 2 with all five morphs combined. The merged pool of males had an average coverage of 78.6X. The initial trimming and mapping of the reads to the genome was done as described below for the whole genome resequencing dataset.

Coverage per position for the pools of females and males was calculated using *SAMtools* (v0.1.19-44428cd) (11) *depth*. We removed positions with: i) mapping quality below 40, ii) with coverage lower than half of the genome-wide average, and iii) with coverage higher than the double of the genome-wide average. We then obtained a ratio, both for males and females, of the average coverage for each chromosome divided by the average coverage of the genome.

#### Quality assessment

We assessed quantitively the completeness of the assembled genome using *BUSCO* (version v3.0.2b) (12,13). Genes contained in *BUSCO* sets for each major lineage are selected from orthologous groups present as single-copy genes in at least 90% of the species. We ran *BUSCO* searches against the tetrapod_odb9 gene dataset.

### Genome annotation

#### Transcriptome sequencing

To obtain empirical information to assist with the genome annotation, we conducted RNA-sequencing (RNA-seq) of five tissues (brain, duodenum, muscle, skin, and testis). The tissues were collected from a second male individual sampled in the same locality as the individual used for the reference genome assembly. After dissection, the tissues were snap frozen in liquid nitrogen and stored at −80°C until RNA extraction. RNA was extracted using the RNeasy Mini Kit with RNase-Free DNase Set (Qiagen). Prior to library construction, RNA integrity, concentration, and quality were assessed using the Agilent Tapestation 2200, Nanodrop, and Qubit RNA (Thermo Fisher Scientific). RNA libraries were prepared using the Illumina TruSeq RNA Sample Preparation kit following the manufacturer’s instructions. The libraries were sequenced on an Illumina Hiseq1500 using 2×125 bp reads. Statistics summarizing the data are given in SI Appendix, Table S3. RNA-seq data from two embryos at 31-somite stage, incubated at 15 or 24°C, were obtained from a previous publication (14) and combined with the newly generated data.

We employed two complementary approaches to assemble *de novo* the transcriptome of the common wall lizard, which we used alongside other sources of information to annotate the reference genome. First, we generated an assembly of normalized reads using the *Trinity* package (v2.2.0) (15) and default settings. This approach was conducted both for each tissue separately and for a combined read file merging all the data. Second, we mapped the RNA-seq reads to the reference genome sequence by means of the *HISAT2* aligner (16) and then used *Cufflinks* (v2.2.1) (17) to obtain genome-guided transcriptome assemblies for each tissue independently. The individual assemblies were merged into a master transcriptome using the tool *Cuffmerge*.

#### Protein database

To provide guidance to the putative coding sequence and structure of annotated features in our reference genome, we obtained high-confidence protein sequence evidence from several sources. First, we extracted 551,705 protein sequences from the Uniprot-Swissprot (18) reference data set (downloaded on 2016-08). This non-redundant collection contains only manually annotated and reviewed proteins. Second, we queried the Uniprot database for protein sequences belonging to the family Lacertidae and extracted a total of 4,794 sequences. Finally, we queried the same Uniprot database and obtained 19,334 protein sequences from the species *Anolis carolinensis*, which is the most comprehensively annotated lizard genome.

#### Repeat masking

We started by creating a species-specific library of repeats by means of the software *RepeatModeler* (http://www.repeatmasker.org/RepeatModeler/). As repeats can be part of actual protein-coding genes, the candidate repeats modelled by *RepeatModeler* were vetted against our proteins set (minus transposons) to exclude any nucleotide motif stemming from low-complexity coding sequences. Based on the created library, the identification of repeat sequences across the genome was performed using *RepeatMasker* (v4.0.3) (19) and *RepeatRunner* (http://www.yandell-lab.org/software/repeatrunner.html). RepeatRunner is a program that integrates *RepeatMasker* results with protein-based information from *BLASTX*. It improves the efficacy of repeat identification by identifying highly divergent repeats, or portions of repeats. It also helps identifying divergent protein coding portions of retro-elements and retro-viruses.

#### Ab-initio training

We opted for an *ab-initio* based annotation strategy combining gene predictions with the available evidence data. The use of multiple gene finders in general improves the genome annotations, therefore gene predictions were computed in several complementary ways. We used *GeneMark* as *ab-initio* predictor due to its effective training on fungal and eukaryote genomes (20,21). We used the algorithm *GeneMark-ES_ET* (v4.3), which integrates information from mapped RNA-seq into the training process. Additionally, we used *AUGUSTUS* (v2.7) (22) and *SNAP* (23). Both algorithms were trained by means of a profile created by running the pipeline *MAKER* (v3.00.0) (24) one time based on the protein evidence described above and the RNA-seq transcript information generated for several tissues. From this gene build, we created a training gene set by selecting the best gene models based on the following criteria: 1) the genes had to be complete (i.e. start/stop codons mandatory); 2) no similarity over 85% was allowed among genes; 3) AED scores (Annotation Edit Distance) had to be lower than 0.1; and 4) genes had to be at a minimum distance of 1,000 bp from each other. In total, 2,828 genes were selected for the *AUGUSTUS* training process.

#### Gene build

High-confidence gene models were computed using the *MAKER* software by combining evidence-based data (protein homology, transcripts, and repeats) with *ab-initio* profiles. An evidence-guided build was computed by allowing the *MAKER* software to construct gene models directly from both aligned transcript sequences and reference proteins. Evidence builds generally closely reflect the information provided by the available sequence data and try to synthesize consensus transcript structures. However, this approach is vulnerable to missing or incomplete sequence material, as well as the noise level into the transcriptome data.

The *ab-initio* evidence driven build was based on the initial evidence annotation by using *MAKER* alignments, together with specifically trained *AUGUSTUS, SNAP* and *GeneMark ab-initio* profiles. The aim of this approach was to perform a second run with all gene models and to replace any gene locus where a longer putative CDS could be predicted by the gene finder or fill in gene predictions where sufficient evidence was lacking from the construction of evidence models.

Statistical evaluation of the final annotation was performed with an in-house (SciLife) Perl script. To improve the annotation, we also ran an in-house script to improve the open reading frame (ORF) start and end, since we noticed that a non-negligible proportion of genes were fragmented and missing from the BUSCO post-annotation validation

#### ncRNA annotation

In addition to protein-coding genes, we also performed an annotation of ncRNA. tRNA were predicted through tRNAscan (v1.3.1) (25). For annotation of other types of broadly conserved ncRNA, we used as the main source of information the RNA family database *Rfam* (v11) (26). *Rfam* provides curated co-variance models, which can be used together with the *Infernal* package (27) to predict ncRNAs in genomic sequences. The set of co-variance profiles was limited to only include broadly conserved, eukaryotic ncRNA families.

#### Functional annotation and gene name inference

Functional inference for genes and transcripts was performed using the translated CDS features of each coding transcript. Each predicted protein sequence was blasted against the Uniprot-Swissprot reference data set in order to retrieve the gene name and the protein function. Each predicted sequence was also blasted against *InterProscan* (v5.7-48) (28) in order to retrieve *Interpro* (29), *Pfam* (30), *GO* (31), *MetaCyc* (32), *KEGG* (33) and *Reactome* (34) information. Finally, using the output from those analyses and the *ANNIE* annotation tool (http://genomeannotation.github.io/annie), we extracted and reconciled relevant meta data into predictions for canonical protein names and functional predictions.

### Whole-genome resequencing of color morphs

#### Sampling strategy

Genome-wide polymorphism data was obtained using whole-genome resequencing of DNA pools. We sampled male individuals of the five color morphs at two localities in the eastern Pyrenees (Angostrina [42°28’N, 1°57’E] and Tor de Querol [42°27’N, 1°53’E]; SI Appendix, Table S4). For each lizard we removed the tip of the tail for genomic DNA extraction, which was performed using the EasySpin kit (Citomed) followed by a RNAse A treatment. DNA quality and concentration were assessed using a NanoDrop spectrophotometer, agarose gels, and the Qubit dsDNA BR Assay Kit (Thermo Fisher Scientific). Individuals were pooled by morph and locality in equimolar concentrations for sequencing. The number of individuals included in each pool varied from 9 to 21.

#### Whole-genome resequencing and quality control

Each pool was sequenced to an effective coverage of 15-18-fold using 2×125 bp reads on an Illumina Hi-seq 1500 (SI Appendix, Table S4). The read data files were initially checked for sequence quality statistics with *FastQC* (v1.7.119) (http://www.bioinformatics.babraham.ac.uk/projects/fastqc). Next, we trimmed the reads to remove adapters, low-quality bases, and flanking regions of each read with *Trimmomatic* (v0.35) (35) using the following parameters: *TRAILING*=15, *SLIDINGWINDOW*=4:20, *MINLEN*=30. Trimmed sequence files were reanalyzed with *FastQC* before proceeding with further analyses.

#### Read mapping and SNP calling

Reads were mapped to our *de novo* reference genome assembly (PodMur1.0) using *bwa-mem* with default settings. Alignment and coverage statistics (SI Appendix, Table S4) were calculated with *SAMtools* and custom scripts. To reduce noise in downstream analyses, we eliminated reads with mapping quality lower than 40 and with overlapping portions between forward and reverse read pairs. This prevented cases in which an allele from an overlapped portion of the same molecule could be considered twice.

These filtered mapped reads were used for variant calling using *FreeBayes* (v1.0.2-33-gdbb6160) (36). *FreeBayes* is a variant detection method that, instead of a traditional alignment-based single-position approach, uses a Bayesian framework to reconstruct haplotypes and call small variants such as SNPs, indels, and other complex variants. *FreeBayes* was run with the 10 pools from both localities using default parameters and setting the ploidy to twice the number of diploid individuals for each pool. Compound variants were decomposed into single variants using the program *decompose-blocksub* from the *vt* tool set (37).

### Population genomics

#### Genetic structure

To infer patterns of population structure among sampled localities and among color morphs we constructed a neighbor-joining tree. We excluded positions with less than 15 reads in any of the pools, followed by a random down sampling of reads in all positions to a maximum coverage of 15X. These filtering steps ensure that low coverage positions are not incorporated in the analysis and that differences in sequencing depth among pools are not biasing estimates of genetic differentiation. These steps were performed using *SAMtools* and *PoPoolation2* (38). From the remaining set of variants, we randomly chose 250,000 SNPs to calculate Nei’s standard genetic distance (39) between each pair of pools based on allele frequencies estimated from allele counts. We then used the distance matrix as input to generate a Neighbor-Joining tree in the *neighbor* program in *PHYLIP* (v3.696) (40).

#### Nucleotide variation and differentiation

To compare levels and patterns of genetic diversity between localities and morphs, we summarized the allele frequency spectrum using Tajima’s *D* (41) and genetic diversity using nucleotide diversity (π) (42). Genetic differentiation was summarized using *F*_*ST*_. As before, reads with a mapping quality lower than 40 and bases with a sequencing quality lower than 20 were not considered. Both π and Tajima’s *D* were calculated in non-overlapping windows of 10,000 bp using *PoPoolation* (43). The same window strategy was used for *F*_*ST*_ calculated by means of *PoPoolation2*. We restricted this analysis to positions with a minimum coverage of 2/3 and a maximum coverage of twice the average coverage of each pool. A genome-wide estimate for each pool was obtained by averaging the values for all windows.

#### Genetic mapping

To test for the association between each color morph and specific genomic regions, we calculated allele frequency differentiation at each previously identified SNP between each relevant comparison. We filtered out SNPs that had coverage higher than two times the average coverage of each pool in at least one of the pools and restricted the minimum coverage to 5x (SNPs with less than this coverage in at least one of the pools were removed). The *PoPoolation2* software was then used to calculate pairwise allele frequency differences between all the pools in each population, for each of these main contrasts based on the phenotypic data:

1. Presence or absence of orange pigmentation: pure orange and mosaic morphs vs. pure white and pure yellow morphs
2. Presence or absence of yellow pigmentation: pure yellow and mosaic yellow morphs vs. white and mosaic-white morphs. For this contrast we removed the pure orange pools – this was done to account for the possibility that the stronger orange coloration could be masking the yellow coloration.
3. Mosaic patterning or uniform coloration: mosaic morphs vs. pure orange morphs.

For each SNP we calculated the median value of allele frequency difference (ΔAF) for each relevant comparison. To reduce the possibility of detecting false positive SNPs in association with each phenotype, we excluded positions that were within 5 bp at both directions from an identified indel using *PoPoolation2* and removed positions that had a minor allele frequency lower than 0.1 in the whole dataset. We then used a sliding window approach (20 SNPs and steps of 5 SNPs) to identify regions with consistent differentiation across many SNPs and avoid spurious associations from individual SNPs derived from erroneous read mapping or stochasticity in allele frequency estimates from pool sequencing.

To estimate whether windows displayed higher allele frequency differentiation than expected by chance, we conducted for each contrast 1000 permutation tests in which we randomized the median allele frequency difference but maintained the genomic positions as in original data. We then applied the same sliding approach and recorded the top value for each permutation. Using this approach, we considered a significant allele frequency difference a value in the original data that was higher than the top 1% of the obtained distribution by permutation. A SNP of high delta within the top windows in each contrast was genotyped by Sanger sequencing, including for the mosaic phenotype where no windows had ΔAF values above the significance threshold.

We further conducted a Cochran-Mantel-Haenszel (CMH) test (44,45). CMH is a repeated test of independence, which assesses consistent allele frequency changes across biological or technical replicates. We used the same filtering steps as for the ΔAF analysis (minimum 5x coverage, maximum double coverage, removal of SNPs within 5 bp around indels, removal of SNPs with minor allele frequency below 0.1), and conducted the test in *PoPoolation2*. In order not to violate the assumption of independence of the method, several replicates for each association were performed, and in each replicate a given pool was used only once:

Presence or absence of orange pigmentation:

- orange vs. white + orange-white vs. yellow
- orange vs. white + orange-yellow vs. yellow
- orange-white vs. white + orange-yellow vs. yellow
- orange-white vs. white + orange vs. yellow
- orange-white vs. orange-yellow + orange vs. yellow
- orange-yellow vs. white + orange-white vs. yellow

Presence or absence of yellow pigmentation:

- yellow vs. orange-white + orange-yellow vs. white
- orange-yellow vs. orange-white + yellow vs. white

Mosaic patterning:

- orange vs. orange-white
- orange vs. orange-yellow

To combine the results for each association test, we summarized *p*-values using the Fisher’s combined probability test (46), and performed the same sliding window analysis as for the ΔAF analysis (20 SNPs and steps of 5 SNPs).

### Confirmation of association by individual genotyping

We confirmed the genotype-phenotype associations in individuals from the Pyrenees through individual-based genotyping using Sanger sequencing. We designed primers to amplify a fragment overlapping the regions associated with phenotypes upstream of *SPR* and *BCO2* (SI Appendix, Table S7). Within each fragment, we tested the association using a single variant showing high allele frequency differentiation between morphs (orange vs. non-orange; yellow vs. white). For the orange locus that variant was a 38-bp insertion relative to the reference (chr9:77,999,995-77,999,996 bp), and for the yellow locus it was a SNP (chr15:26,161,682 bp). A significant association between these variants and their respective phenotypes was tested using the R package *SNPassoc*) v1.9-2) (47). These same variants were genotyped in additional individuals belonging to other sub-lineages of the common wall lizard.

### Nanopore sequencing

#### DNA-preparation

To investigate structural variation around the candidate loci, we sampled skeletal muscle from an orange-white individual (from locality 2) and extracted high-molecular weight DNA using a modified phenol-chloroform extraction (https://www.protocols.io/view/ultra-long-read-sequencing-protocol-for-rad004-mrxc57n). We grinded the tissue with a pre-frozen mortar and pestle over dry ice and incubated the tissue for 5 h in TE with proteinase K and RNase A. After digestion, an equal volume of phenol-chloroform-isoamyl alcohol (25:24:1) was added, mixed gently for 10 min and centrifuged at 4,500 rpm for additional 10 min. This step was repeated a second time using an equal volume of chloroform-isoamyl alcohol (24:1). The aqueous phase was mixed with 1:10 volume of ammonium acetate (5M) and 2:1 of 100% ice-cold ethanol and mixed gently. DNA was then physically removed from the solution and placed in a tube with 70% ethanol, which was centrifuged at 10,000 g. The washed DNA pellet was dried and then dissolved in 1x TE buffer. Dissolved DNA was subjected to mechanical shearing to 15 kb and 25 kb fractions using a Megaruptor2 instrument (Diagenode).

#### Oxford Nanopore library generation and sequencing

The sheared DNA-fractions were cleaned up using Ampure XP beads (Beckman Coulter) and the DNA was eluted in nuclease-free water. We produced two Oxford Nanopore libraries for sequencing in a MinION device, starting from 7.6 μg of DNA each (15 kb and 25 kb fractions, separately) using the SQK-LSK108 kit, with DNA-repair and DNA end-prep combined according to the one-pot protocol (https://www.protocols.io/view/one-pot-ligation-protocol-for-oxford-nanopore-libr-k9acz2e). After the combined FFPE DNA repair/end repair step we cleaned the repaired/end-prepped DNA using Ampure XP beads and eluted in nuclease-free water. Adapter ligation, and all following library preparation steps, was carried out using the official protocol (1D Genomic DNA by ligation) from Oxford Nanopore (www.nanoporetech.com). We loaded 0.8 and 2 μg, onto two different FLO-MIN106 (r9.4.1) flow cells for the 15 kb and 25 kb libraries, respectively. Sequencing was run for 48 h and we obtained 2,320,498 sequence reads with an average length of 8,730 bp, resulting in an average genome-wide coverage of 9.7X (84.9% of the total reads mapped against the reference genome). The resulting raw data fast5 files were base-called using *guppy* (v0.3.0) (Oxford Nanopore).

#### Analysis of long read sequencing data

The resulting sequence files were subjected to adapter trimming using *downpore* (v0.2) (J. Teutenberg, downloaded from: https://github.com/jteutenberg/downpore). Trimmed files were mapped to our reference assembly using *NGMLR* (v0.2.6) (48) with default parameters. Since our objective was to investigate structural variation in the regions associated with pigmentation, we retrieved reads that mapped within and around *SPR* and *BCO2*. We manually realigned the reads in BioEdit (v7.2.5) (49) to construct a consensus sequence based on the plurality rule (i.e. for each position we obtained the most frequent state, even if not found in the majority of sequences), using a script by Joseph Hughes (University of Glasgow, downloaded from https://github.com/josephhughes) and aligned the consensus to the reference genome sequence. The results were visualized using *Jalview* (v2.10.4b1) (50).

### Annotation of SNPs and indels

Variant annotation was performed using the genetic variant annotation and effect prediction toolbox *SnpEff* (v4.3) (51). We specifically searched for SNP and indel variants with potential functional significance around the candidate regions and that have the potential to alter protein structure, such as frameshift, nonsynonymous, stop, and splice site mutations.

### Gene expression and allelic imbalance analysis

#### Quantitative real-time PCR analyses of gene expression

Gene expression was quantified using quantitative PCR. We sampled 26 individuals belonging to all five morphs (six white, six orange, five yellow, five orange-white and four orange-yellow) from Llívia (42°27’N, 1°58’E). After dissection, several organs (skin, brain, liver and muscle) were harvested, snap-frozen in liquid nitrogen, and stored at −80°C until RNA extraction. RNA was extracted using the RNeasy Mini Kit (Qiagen). cDNA was generated by reverse transcribing ∼1 μg of RNA using the GRS cDNA Synthesis Kit (GRiSP) following the manufacturer’s protocols. For the analysis on the skin, we focused only on the pure morph animals, to avoid possible variation in expression across skin patches of mosaic animals.

We designed primers located on different exons to minimize amplification from contaminant genomic DNA for the following genes, *SPR, BCO2, PTS*, and *18S* (housekeeping gene used for expression normalization across samples) (SI Appendix, Table S7). Three independent assays were performed for each biological replicate on a CFX96 Touch Real-Time PCR Detection System using iTaq Universal SYBR Green Supermix (Bio-Rad Laboratories). Cq (quantification cycle) values of the three technical replicates were then averaged and the expression of each focal gene for each sample was normalized to the expression of *18S* using a -ΔCq approach (52). Finally, we compared the normalized expression of each gene between color morphs in a pairwise manner using the Mann-Whitney U test. For the test of *PTS* expression on the skin, one individual of the white morph was excluded, as it showed only residual PCR amplification possibly explained by a mutation in the primer binding site.

#### Quantification of allelic imbalance

We also assessed levels of gene expression by analyzing allele imbalance. Since the association does not extend to the coding region in *SPR* and weakly so in *BCO2*, we could not examine a diagnostic variant between pigmentation alleles to quantify allelic expression. Thus, we cannot deduce which specific allele is preferentially expressed at these loci due to the lack of linkage disequilibrium between the non-coding sequences showing an association and SNPs located in the transcript. However, if the color polymorphisms are controlled by cis-regulatory mutations affecting the expression of *SPR* and/or *BCO2*, we expect to see allelic imbalance in expression in individuals that are heterozygous *O/o* and *Y/y*. For all individuals that among our cohort of samples used for the qPCR analysis were heterozygous *O/o* and *Y/y*, we screened part of the coding sequence to detect polymorphisms that could be used to quantify the relative expression of each allele. We then designed primers to amplify small fragments from cDNA overlapping these polymorphisms. Primers were located on exon-exon junctions to minimize amplification from genomic DNA and were carefully placed to avoid overlapping existent polymorphisms (SI Appendix, Table S7).

The allelic expression was quantified by sequencing on a MiSeq (MiSeq v3 600-cycle kit, 2×300 bp reads). Amplification of the cDNA template with 5’ labeled primers and pre-processing of the reads prior to mapping to the genome was done with the same protocol as described in more detail below (“*Amplicon sequencing overlapping regions of association*”). To calculate the relative proportion of alleles expressed in the skin of each individual for each transcript, we counted the number of reads corresponding to the reference and alternative alleles. Finally, allelic imbalance was tested using chi-square tests (a proportion of 1:1 was used as the null hypothesis).

### Amplicon sequencing overlapping regions of association

#### Sampling and PCR amplification

We sequenced amplicons (∼550 bp) overlapping the regions of association for the two loci recovered in the genome-wide association mapping. We sequenced a set of samples (n=48) that included common wall lizard samples from our primary study location in the eastern Pyrenees (n=16) and other *Podarcis* species (n=32) that are known to present ventral color polymorphism (further details can be found on SI Appendix, Table S8). Genomic DNA was extracted from tail-tip samples using the EasySpin kit (Citomed) followed by a RNAse A treatment.

Amplification of the target regions was done via a two-step PCR protocol based on (53). Briefly, a first PCR with 5’-tailed primers served to amplify the target region, followed by a second PCR to attach barcoding sequences to the amplified DNA (SI Appendix, Table S7). The first PCR reaction was prepared with approximately 25 ng DNA, 5 µL 2x Qiagen MasterMix, 0.4 µL of 10 pM of each primer and 3.2 µL PCR-grade water, and was run under the following conditions: 1) an initial denaturing step of 95°C for 15 min; 2) 5 touch-down cycles with 95°C denaturing for 30 s, a 68-64°C annealing temperature touchdown for 30 s and 72°C extension temperature for 45 s; 3) 35 cycles with 95°C denaturing for 30 s, a 64°C annealing step for 30 s and 72°C extension for 45 s; 4) a final extension at 60°C during 20 min. Before conducting the second PCR we purified the resulting DNA with a standard bead cleaning protocol (a 0.7:1 bead-to-sample volume ratio was used). We set up the second (barcoding) PCR reaction using 2 µL of purified PCR product, 5 µL 2x Qiagen MasterMix, 1 µL of a mix of individually labeled primers with P5/P7 binding sites and 1 µL of PCR-grade water. The following program was used for the barcoding PCR: 1) an initial denaturing step of 95°C for 15 min; 2) 10 cycles with 95°C denaturing for 5 s, a 55°C annealing temperature step for 20 s and a 72°C extension for 45 s; 3) a final extension at 60°C during 20 min. After a second bead cleaning step, all samples were pooled at equimolar concentrations. The pooled library was loaded onto a MiSeq (MiSeq v3 600-cycle kit, 2×300 bp reads) at a concentration of 13.5 pM.

#### Sequence processing and haplotype inference

After demultiplexing the raw reads, we removed low quality reads, low quality bases, and primer sequences with *Trimmomatic* using the following parameters: *TRAILING*:15, *HEADCROP*:22, *SLIDINGWINDOW*:4:20, *MINLEN*:30. Since the sequence length of paired-end reads combined was larger than the fragment size, paired-end reads were merged into a single sequence with the *fastq-join* utility (54). To filter out non-target amplified DNA, we mapped the new merged reads using *bwa-mem* with default parameters to a small reference sequence containing the two target regions and approximately 100 bp around each locus. The reads that mapped to each of the two regions were then reconverted into separate fastq files and the fastqs were randomly down-sampled to 10% of the total number of reads to decrease computational times in subsequent analyses. Both approaches were carried out with *picard* (http://broadinstitute.github.io/picard).

Given that the divergence between haplotypes was extremely high (see main manuscript), traditional mappers and SNP callers produced erroneous datasets. Therefore, to obtain variant and haplotype information from each individual/locus we used a custom python script (available upon request). Briefly, the script begins by defining an average sequence similarity threshold between a subset of all unique reads present in the dataset (reads are mapped pairwise using dynamic programming; 10% of reads were used by default). If the average sequence similarity within the subset exceeded 99.5% (equivalent to only one or two mutations in a 500 bp amplicon), this was used as the reference value. After defining the threshold, all unique reads from the dataset appearing at least five times were sorted by frequency of occurrence. Starting from the most frequently occurring unique read (which is considered one of the haplotypes), the program sequentially maps the following unique reads to the first read using dynamic programming with two possible outcomes: 1) if the sequence similarity between the two reads exceeds the average threshold that was previously estimated (indicating a very similar sequence), the second read is discarded and assigned to the haplotype; 2) if the sequence similarity between the two reads is lower than the average threshold that was previously estimated (indicating a divergent sequence), the second read is kept as a new haplotype for the following pairwise comparisons. This approach led to most samples either having one or two representative sequences. For ambiguous samples with three or more estimated haplotypes, we manually curated the result by analyzing the mapping files using *IGV2*.*4*.*10* (http://www.broadinstitute.org/igv). Finally, we manually realigned the sequences in *BioEdit* (v7.2.5) to build our final dataset.

#### Intra-and inter-specific haplotype networks

We used *popart* (v1.7) (55) to construct and plot median-joining haplotype networks. To explore patterns of sequence evolution in a broader phylogenetic context, we constructed neighbor-net networks using *SplitsTree* (v4.14.2) (56) using an extended dataset with additional sequences from other *Podarcis* species. A neighbor-net approach was used because (unlike traditional phylogenetic trees) it considers processes like recombination and hybridization. To simplify the visual representation of the phylogenetic networks we used the following criteria to subsample the alignment dataset: 1) since the initial within-population median-joining network analysis detected 2-3 highly divergent haplogroups, we randomly selected one sequence from each haplogroup to represent *P*. *muralis*; 2) from each of the other species (for each tree, orange locus and yellow locus) we randomly selected two sequences from individuals with the relevant pigmentation (orange or yellow) and two sequences from individuals without the pigment. We checked for potential biases introduced by this subsampling by repeating the analysis with other sequences multiple times, and confirmed that the general pattern remained qualitatively unchanged. Indels were not considered in both analyses.

#### Haplotype divergence

We estimated pairwise sequence divergence between haplotypes of the *SPR* and *BCO2* amplicons within *P*. *muralis* and between *P*. *muralis* and six additional *Podarcis* species. In addition, we performed the same calculations for 31 amplicons randomly distributed in the genome (SI Appendix, Table S9). These amplicons ranged from 255 bp to 667 bp (average = 410 bp) (57-62). Part of the data was obtained from a previous publication (63), but we PCR and Sanger sequenced some amplicons to complete the data for the six additional species. PCR protocols were carried out according to (63).

## SUPPLEMENTARY FIGURES AND TABLES

**Figure S1.**
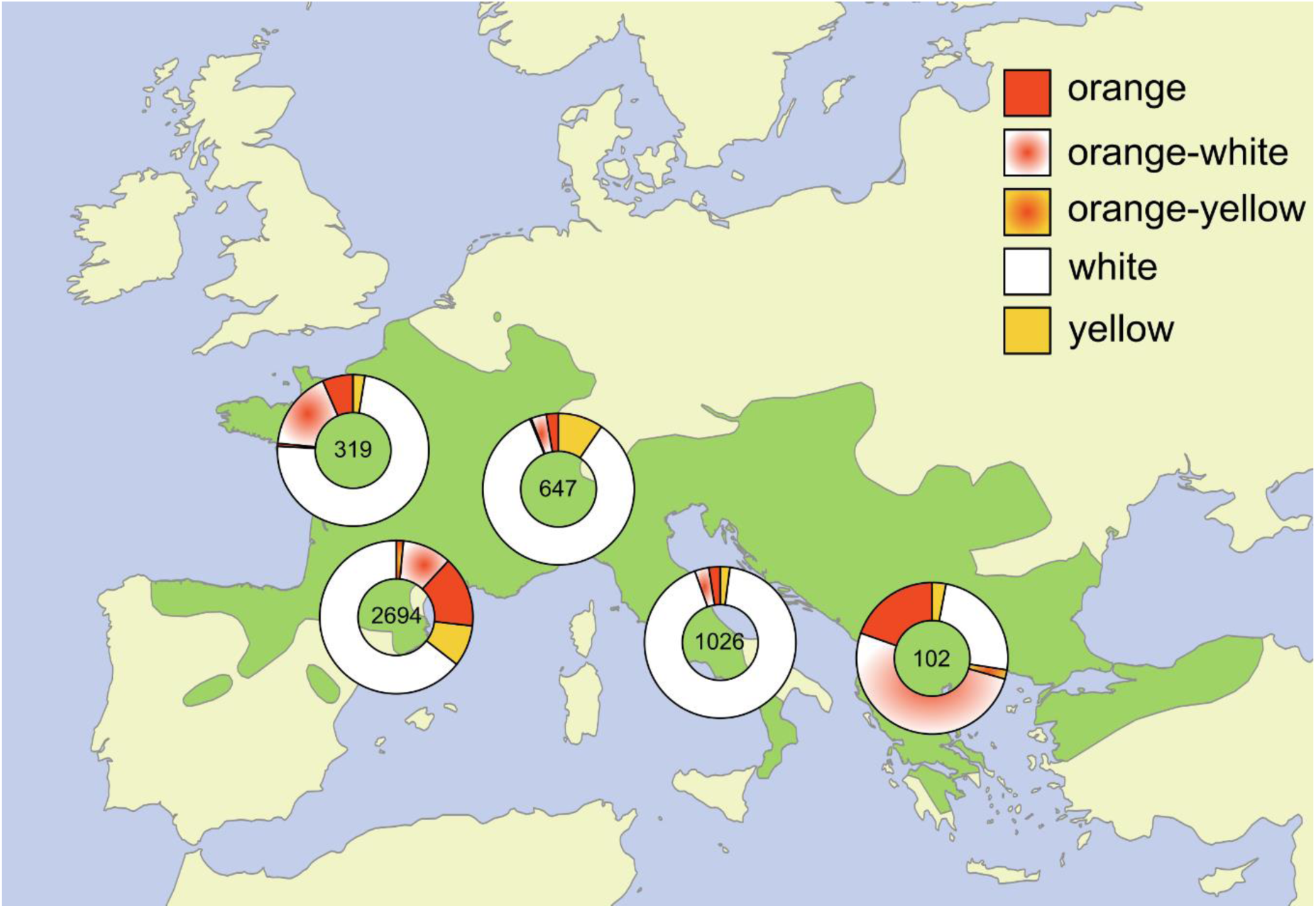
Distribution of the common wall lizard (*Podarcis muralis*) in Europe (green shading), with information on morph frequencies in several locations, showing the wide distribution of this polymorphism across the species’ distribution. Morph frequencies vary at a regional level as well as on a micro-geographic scale. The number of individuals sampled in each locality is indicated.

**Figure S2.**
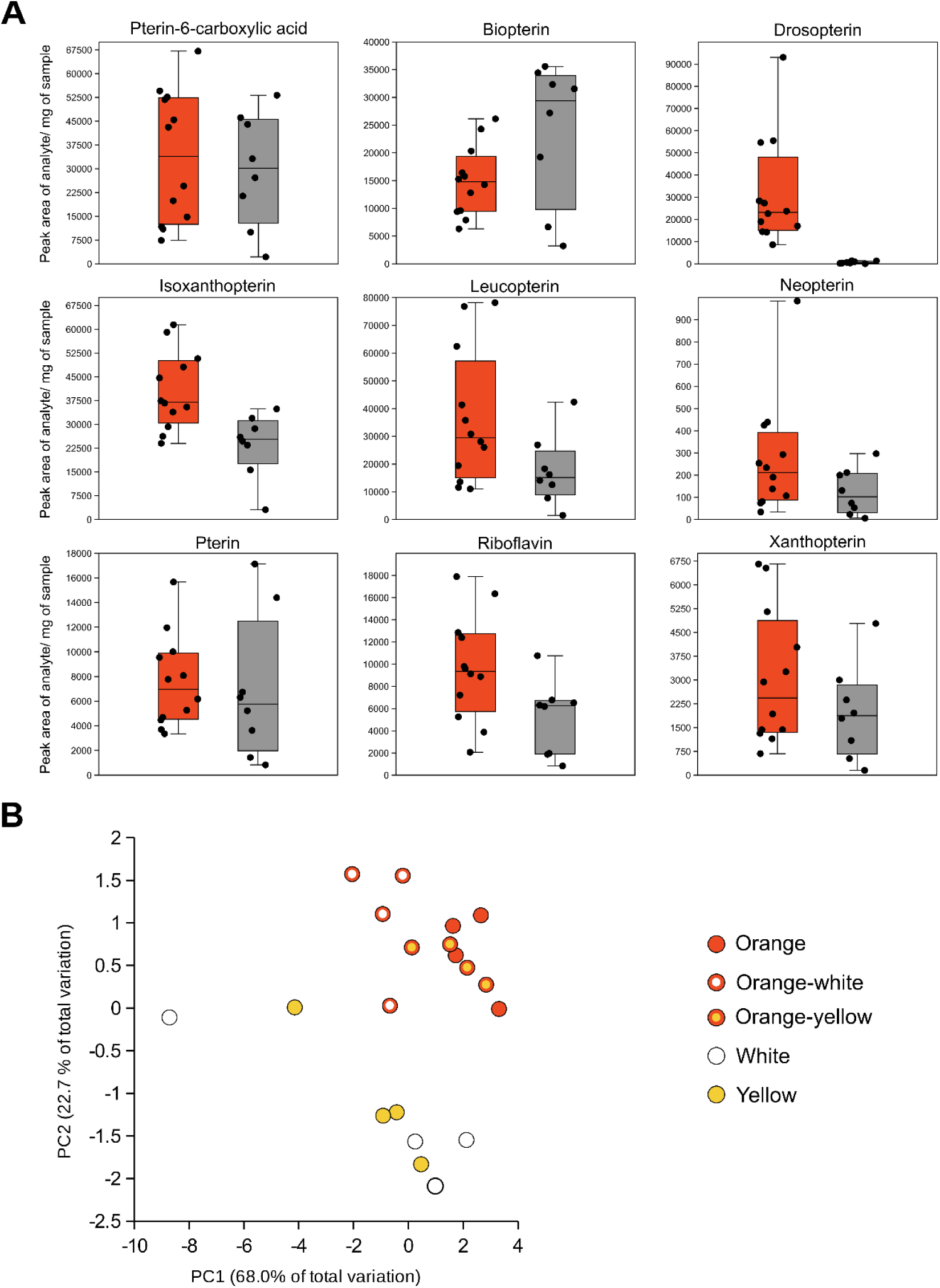
Quantification of pterin compounds (HPLC-MS/MS) in animals expressing orange, yellow and white coloration. **(A)** Results for individual pterin compounds. Orange, orange-white, and orange-yellow individuals are collapsed in a single group, and the same for individuals of the white and yellow morphs. **(B)** Principal component analysis using all pterins as variables, indicating that individuals expressing orange coloration (orange and both mosaic morphs) differ from white and yellow individuals in their overall pterin profile.

**Figure S3.**
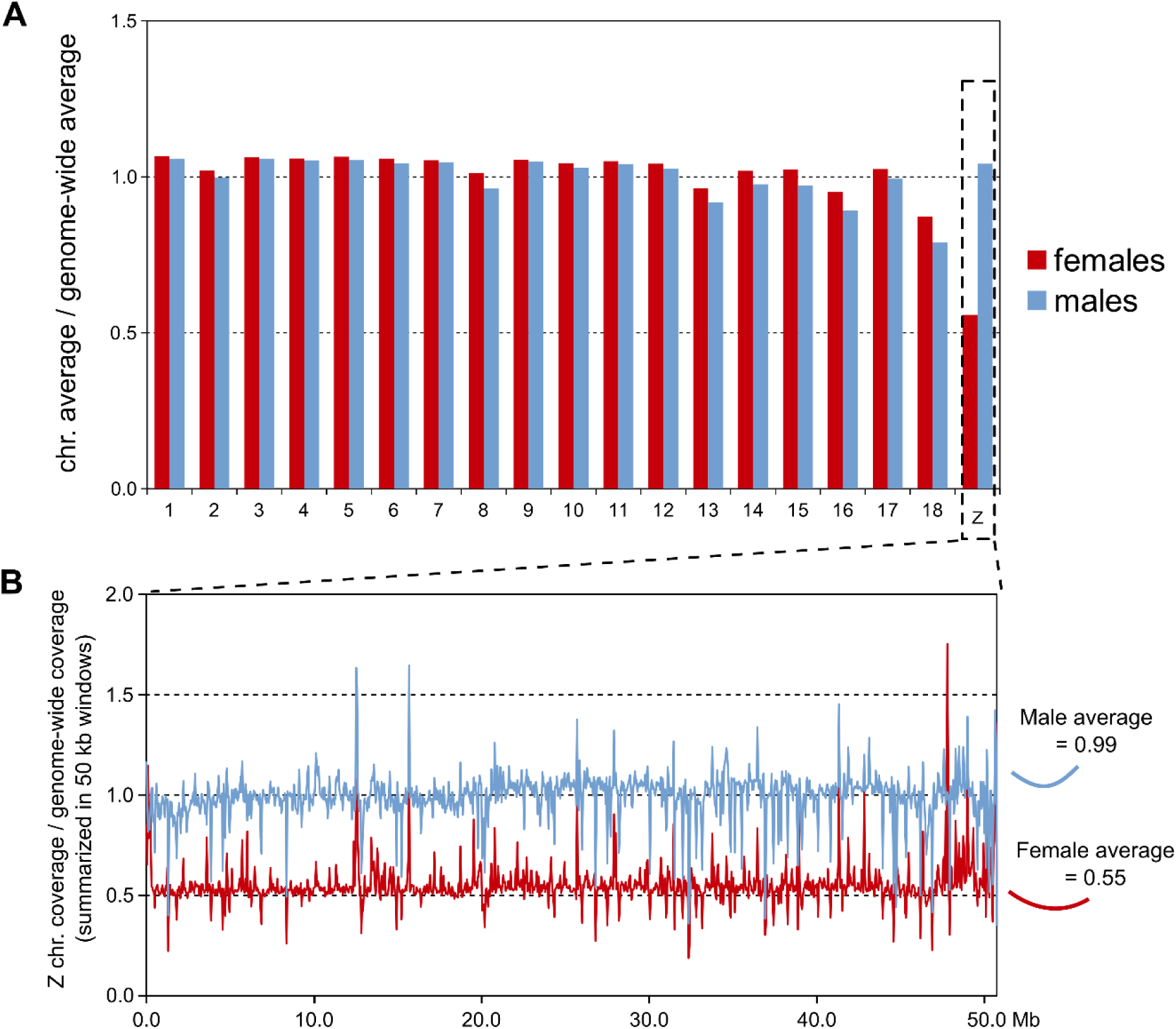
Identification of the Z chromosome in the common wall lizard *de novo* reference assembly using pool-sequencing data of females (ZW) and males (ZZ). **(A)** Ratio between the average coverage of each of the 19 chromosomes and the average genome-wide coverage. The average was calculated per position. Consistent with a ZW sex determination, coverage for one scaffold, that we named Z, was reduced by half in females when compared to males. The ratio between the average number of reads along the candidate Z chromosome and the average number of reads genome-wide. The average coverage was summarized in 50 kb windows.

**Figure S4.**
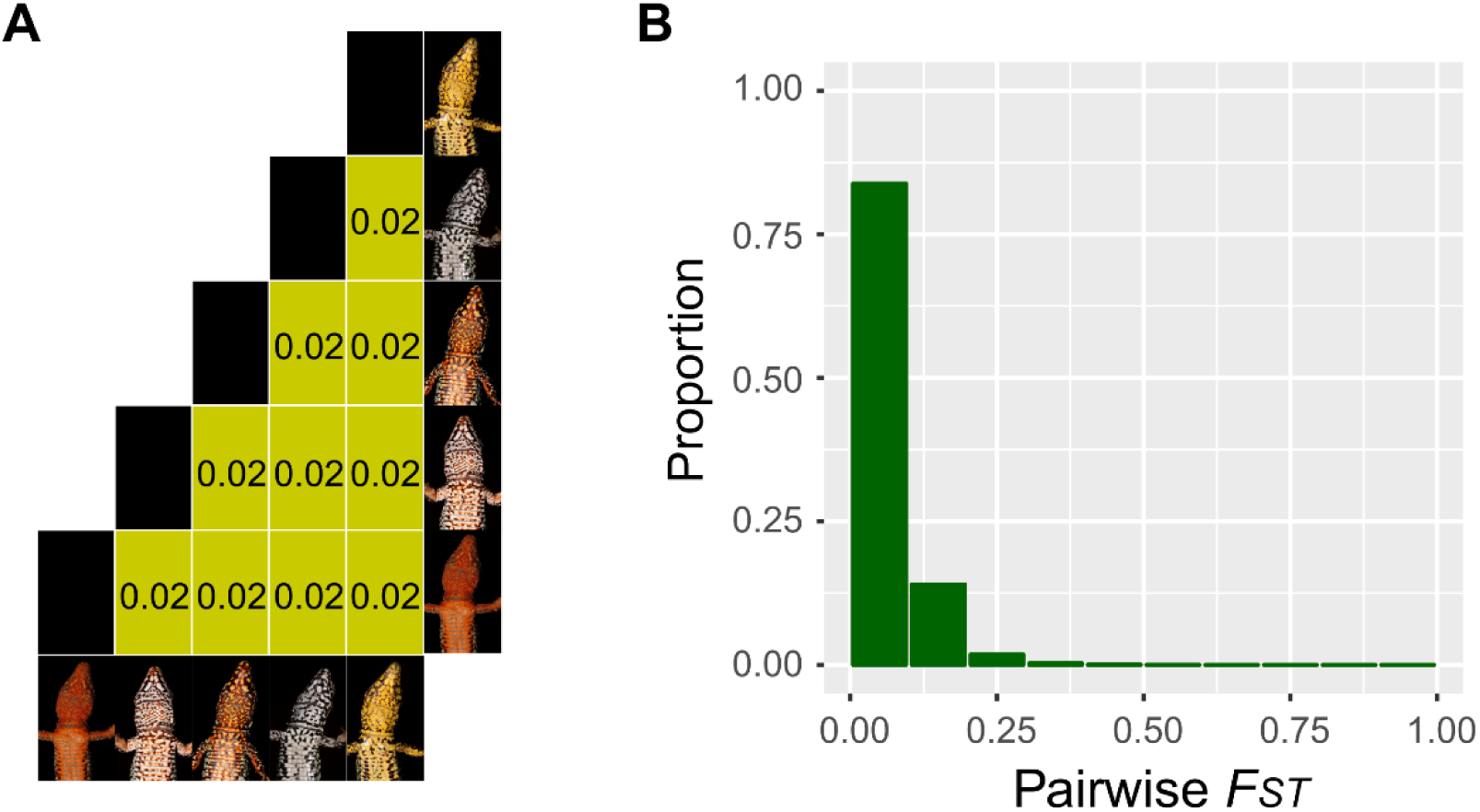
Genetic differentiation among the five color morphs of the common wall lizard calculated across the genome in non-overlapping 10 kb windows. **(A)** Pairwise *F*_*ST*_ values between the five color morphs. **(B)** Distribution of pairwise *F*_*ST*_ values between morphs in the population.

**Figure S5.**
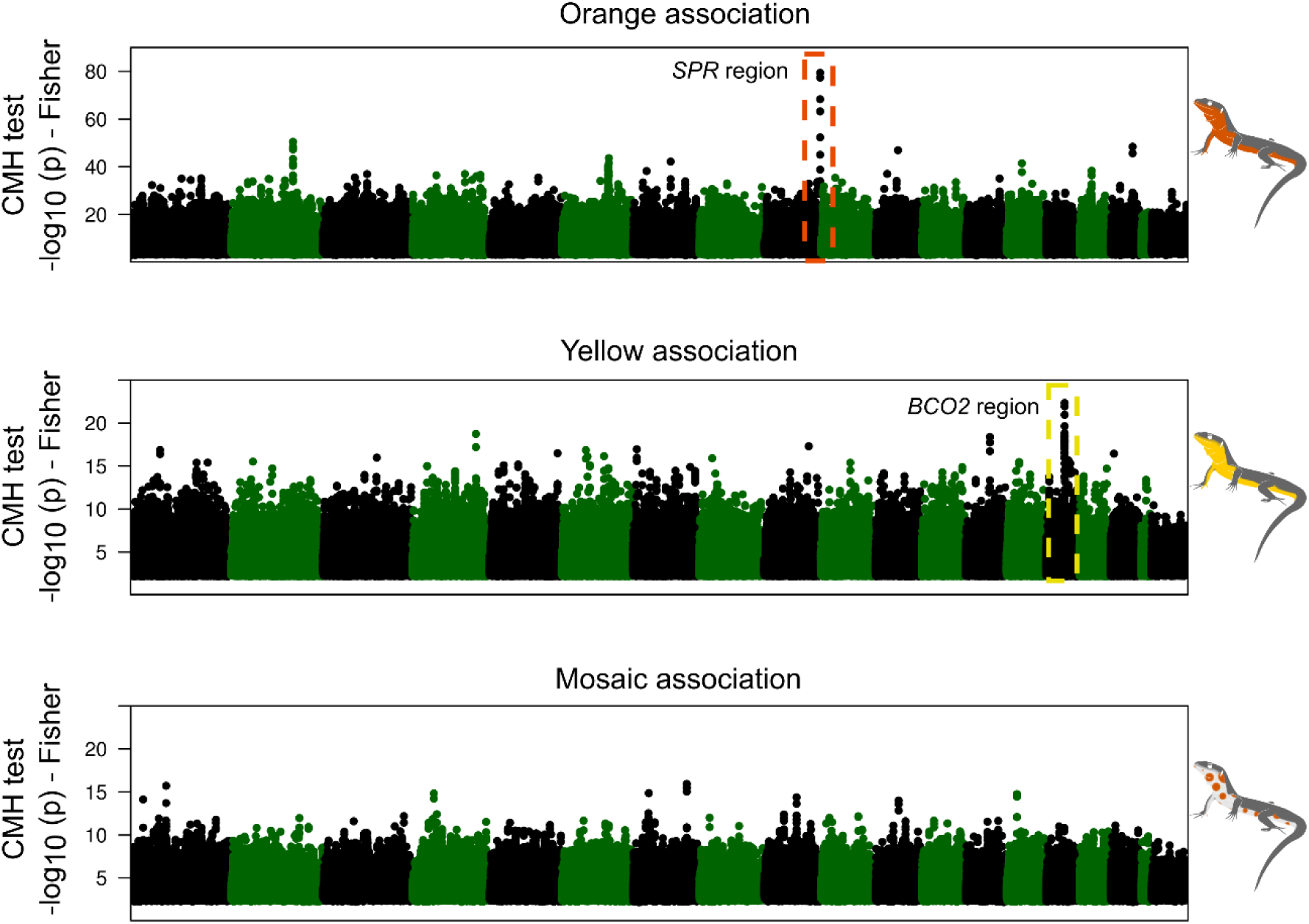
Genome-wide results from a Cochran-Mantel-Haenszel (CMH) test of association. Results were summarized using a sliding-window approach. Each dot represents a 20-SNP window with a step of five SNPs.

**Figure S6.**
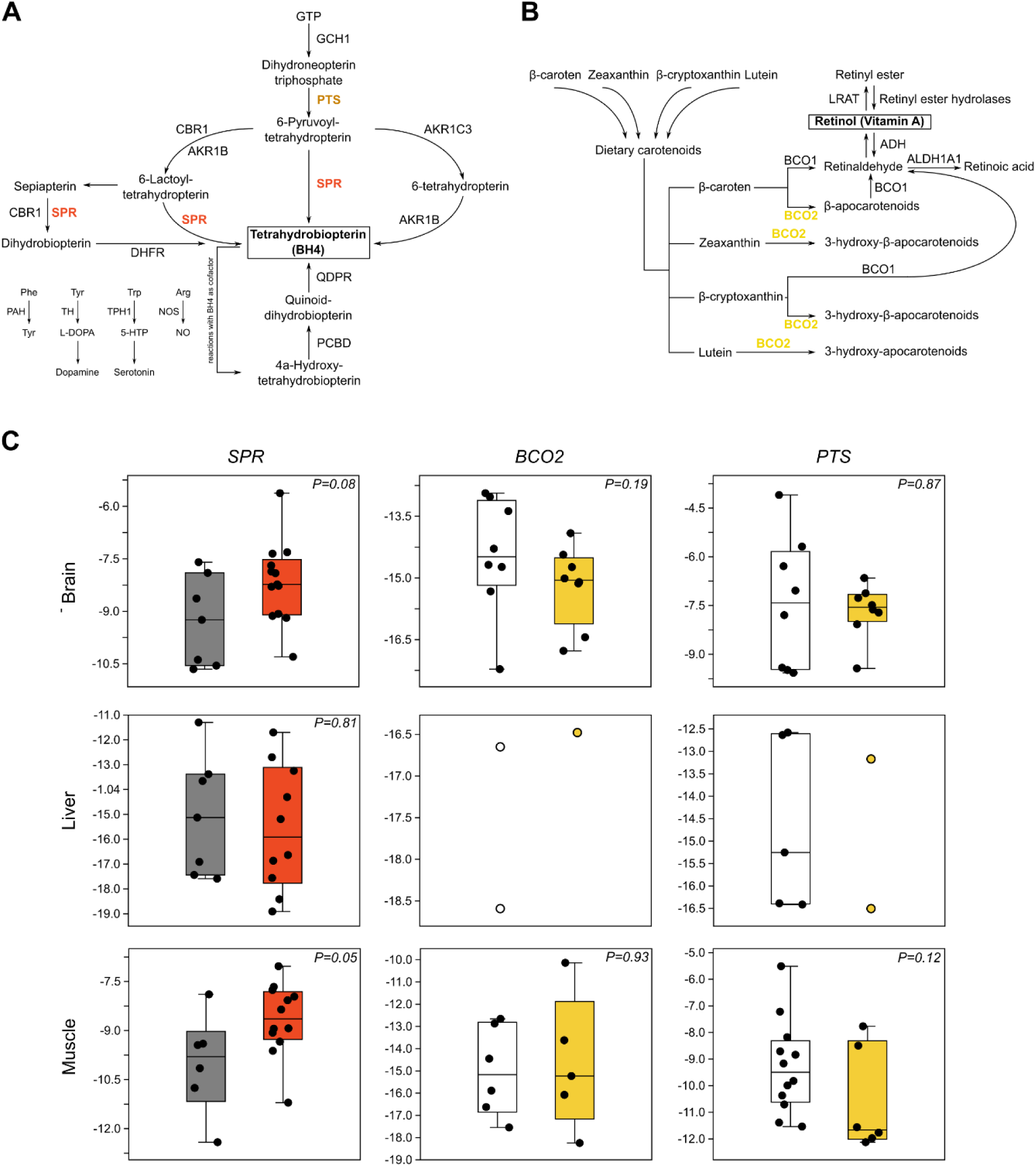
Pterin and carotenoid genes near the regions of maximum association are involved in vital metabolic processes. **(A)** Simplified tetrahydrobiopterin (BH4) biosynthesis pathway, evidencing the crucial roles of SPR and PTS in many of the major conversion steps, in particular of the main *de novo* synthesis pathway. BH4 is an essential co-factor in several enzymatic reactions, notably in those producing neurotransmitters (dopamine, serotonin) and nitric oxide, as well as the metabolism of phenylalanine. **(B)** Simplified view of the main reactions involved in the metabolism of precursor dietary carotenoids, with emphasis on the role of *BCO2*. **(C)** Gene expression patterns of *SPR, BCO2* and *PTS* in brain, liver and muscle, based on qPCR, between animals with orange/non-orange pigmentation and white/yellow pigmentation (y axis shows -ΔCq values, normalized to 18S gene expression). Many samples failed amplification for *BCO2* and *PTS* in the liver, which is likely due to low expression levels of these genes or alternative splice variants specific to this organ. It can also be explained by degradation of the mRNA since RNA integrity numbers for liver were on average lower than that for the other tissues.

**Figure S7.**
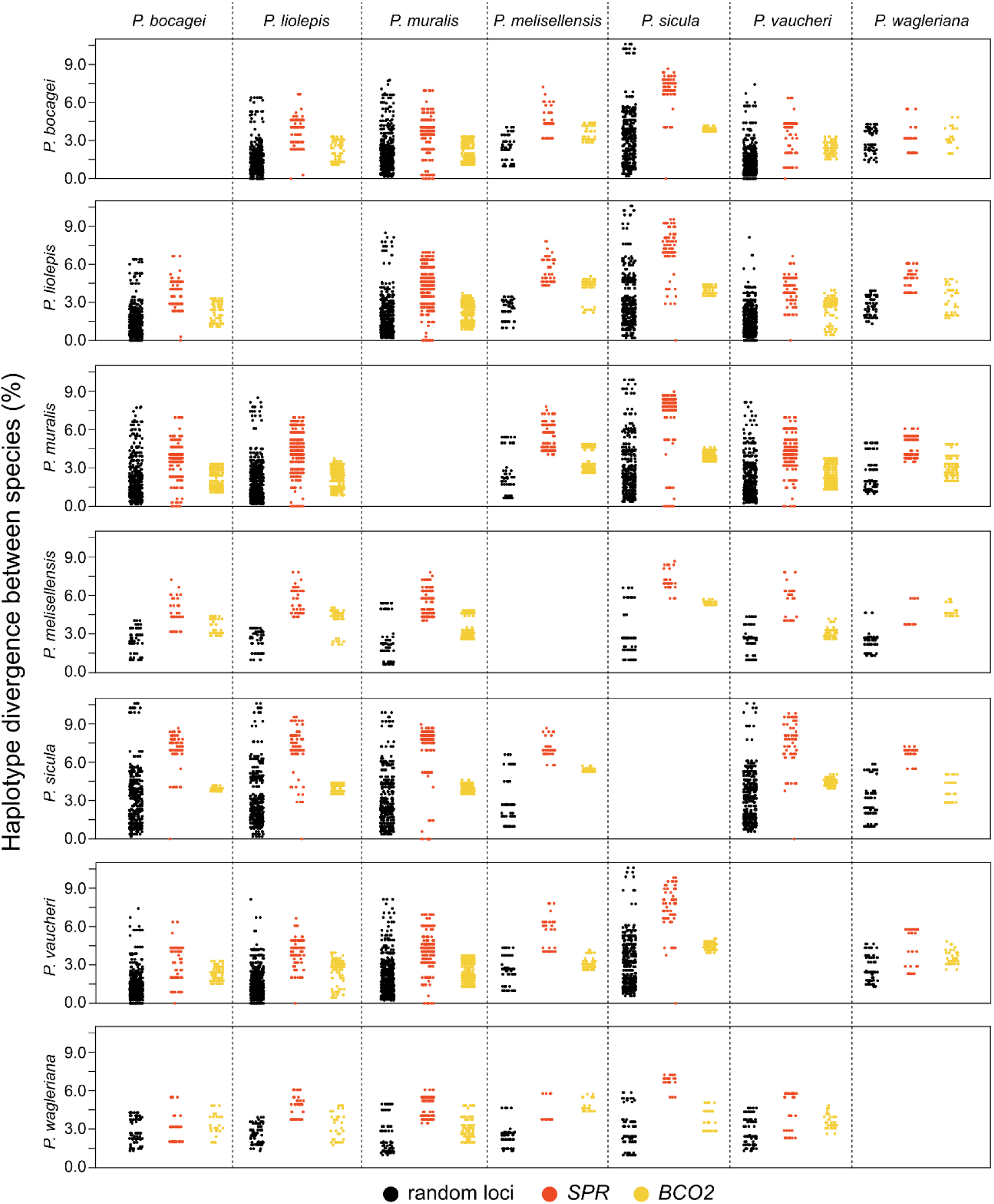
Pairwise nucleotide differences between haplotypes from seven *Podarcis* species for *SPR* (orange), *BCO2* (yellow), and 31 random loci (black).

**Table S1.** Summary statistics for our *de novo* common wall lizard (*Podarcis muralis*) reference genome assembly and annotation (PodMur 1.0)

|  | **PodMur1.0** |
| --- | --- |
| **Genome assembly** |  |
| Contig N50 (kb) | 714.6 |
| Scaffold N50 (Mb) | 92.4 |
| Assembly size (Gb) | 1.51 |
| Number of scaffolds (>1kb) | 2,122 |
| Number of scaffolds (>10kb) | 1,566 |
| Largest scaffold (Mb) | 140.2 |
| GC content (%) | 44.1 |
| Complete BUSCOs reference (%) | 93.2 |
| Fragmented BUSCOs reference (%) | 3.8 |
| Missing BUSCOs reference (%) | 3.0 |
| **Genome annotation** |  |
| Protein-coding genes | 24,656 |
| Complete BUSCOs annotation (%) | 77.2 |
| Fragmented BUSCOs annotation (%) | 11.4 |
| Missing BUSCOs annotation (%) | 11.4 |

**Table S2.** Size of the 19 largest scaffolds of *de novo* reference assembly of the Common wall <u>lizard genome (PodMur1.0).</u>

| Chromosome | Size (bp) |
| --- | --- |
| 1 | 140,209,921 |
| 2 | 128,038,163 |
| 3 | 124,954,059 |
| 4 | 107,526,675 |
| 5 | 100,393,251 |
| 6 | 99,678,440 |
| 7 | 92,398,148 |
| 8 | 90,342,841 |
| 9 | 79,135,106 |
| 10 | 76,280,819 |
| 11 | 65,231,656 |
| 12 | 60,949,594 |
| 13 | 56,546,671 |
| 14 | 54,988,359 |
| 15 | 46,441,056 |
| 16 | 43,659,005 |
| 17 | 42,925,489 |
| 18 | 14,237,138 |
| Z | 50,871,175 |
| Unplaced scaffolds | 36,195,651 |
| **Total** | **1,511,003,217** |

**Table S3.** Summary statistics of RNA sequencing data of several tissues for genome annotation

| **Tissue** | **Number of reads** | **Number of mapped reads** | **Estimated number of transcripts<sup>1</sup>** |
| --- | --- | --- | --- |
| Brain | 230,727,821 | 207,227,126 (89.8%) | 35,281 |
| Duodenum | 129,921,036 | 109,579,178 (84.3%) | 32,159 |
| Embryo (15°C) | 76,264,214 | 70,123,642 (92.0%) | 34,820 |
| Embryo (24 °C) | 69,718,379 | 63,868,404 (91.6%) | 29,689 |
| Muscle | 140,540,165 | 129,421,702 (92.1%) | 20,923 |
| Skin | 138,343,774 | 111,773,184 (80.8%) | 21,512 |
| Testis | 220,430,106 | 197,029,219 (89.4%) | 59,277 |

<sup>1</sup>Estimated using *Cufflinks*

**Table S4.** Summary statistics of the whole genome resequencing dataset

| Population | Locality<sup>1</sup> | Morph | Number of individuals in pool | Number of reads | % of reads mapping<sup>2</sup> | % of reads MQ>=40 | Mean depth of coverage |
| --- | --- | --- | --- | --- | --- | --- | --- |
| Pyrenees | 1 | Orange | 14 | 228,390,110 | 98.2 (91.9) | 82.8 | 17.1 |
| Pyrenees | 1 | Orange-white | 14 | 210,469,900 | 98.6 (92.3) | 83.0 | 15.7 |
| Pyrenees | 1 | Orange-yellow | 15 | 203,437,559 | 98.6 (92.3) | 83.1 | 15.2 |
| Pyrenees | 1 | White | 15 | 215,363,776 | 98.6 (92.4) | 83.2 | 16.1 |
| Pyrenees | 1 | Yellow | 15 | 199,797,350 | 98.4 (92.1) | 82.9 | 14.9 |
| Pyrenees | 2 | Orange | 20 | 216,047,786 | 98.3 (91.8) | 83.1 | 16.3 |
| Pyrenees | 2 | Orange-white | 11 | 201,814,386 | 98.5 (92.0) | 83.4 | 15.3 |
| Pyrenees | 2 | Orange-yellow | 9 | 196,309,282 | 98.5 (92.1) | 83.3 | 14.8 |
| Pyrenees | 2 | White | 21 | 200,992,353 | 98.5 (91.8) | 83.2 | 15.1 |
| Pyrenees | 2 | Yellow | 20 | 240,708,482 | 98.5 (92.0) | 83.4 | 18.2 |
| Italy | - | NA | 20 | 321,742,840 | 98.1 (84.9) | 77.1 | 23.5 |
| Pyrenees (male pools merged) | 2 | NA | 81 | 1,057,458,695 | 98.5 (92.9) | 83.1 | 78.6 |
| Pyrenees (female pool) | 2 | NA | 24 | 755,245,078 | 97.9 (92.2) | 83.0 | 65.5 |

<sup>1</sup>Locality 1 - Angostrina (42°28'N, 1°57'E); Locality 2 - Tor de Querol (42°27'N, 1°53'E).
<sup>2</sup>The percentage of properly paired reads are indicated in parenthesis.

**Table S5.**
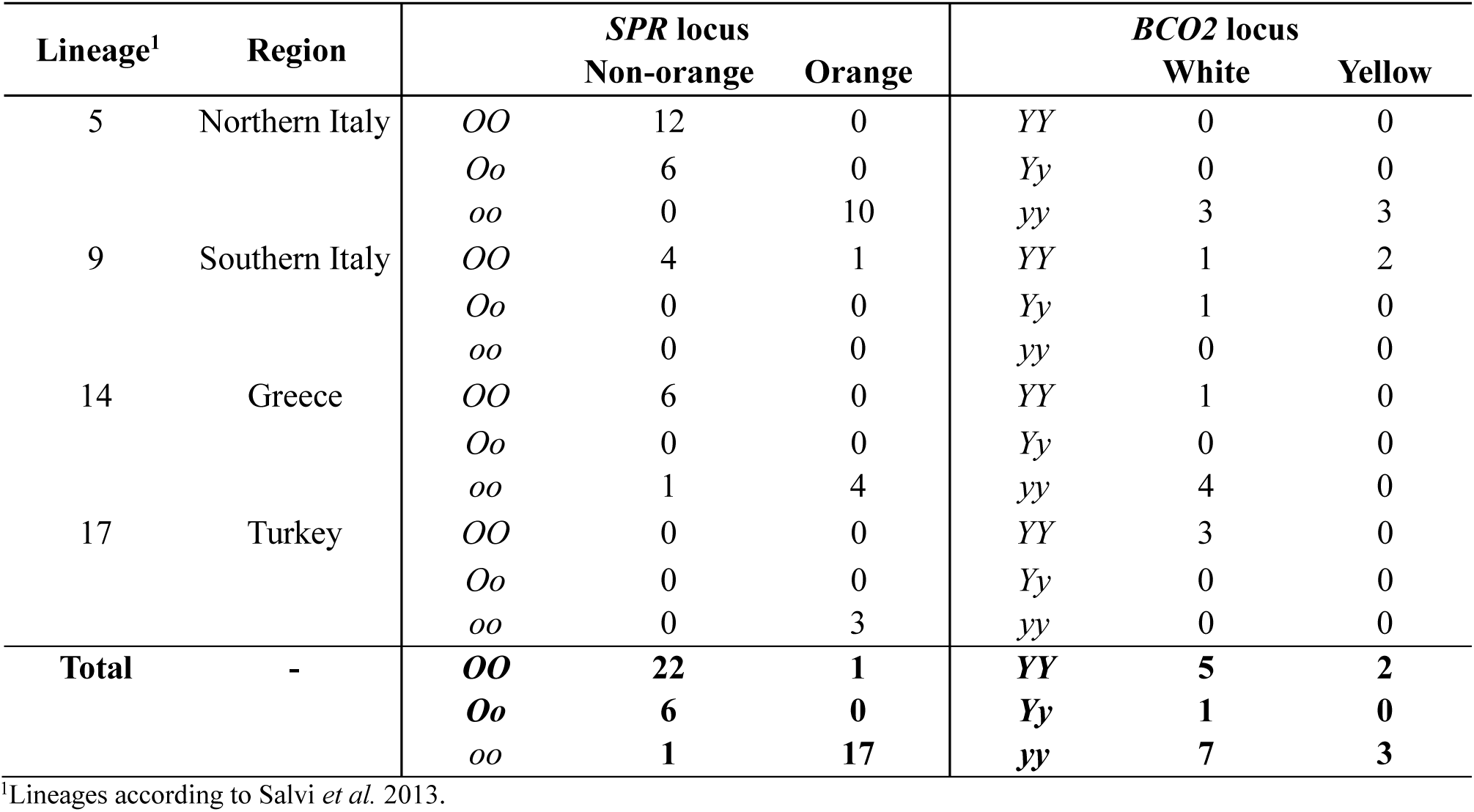
Genotyping results for the *SPR* and *BCO2* loci in other sub-lineages of the common wall lizard

**Table S6.** SRM conditions used for LC-MS/MS determination of the pterin compounds

| **Compound** | **Molecular weight** | **Precursor ion** | **Product ion** | **Fragmentor (V)** | **Collision energy (V)** |
| --- | --- | --- | --- | --- | --- |
| Riboflavin | 376.4 | 377.4 | 243.3 | 135 | 30 |
| L-Sepiapterin | 237.2 | 238.2 | 192.1 | 120 | 15 |
| Pterin | 163.1 | 164.1 | 119.1 | 100 | 20 |
| Pterin-6-carboxylic acid | 209.2 | 208.1 | 162.1 | 100 | 15 |
| 6-Biopterin | 237.1 | 238.1 | 178.1 | 115 | 17 |
| Isoxanthopterin | 237.1 | 238.1 | 178.1 | 115 | 20 |
| Xanthopterin | 179.1 | 180.1 | 135.0 | 125 | 20 |
| Leucopterin | 195.1 | 196.1 | 140.1 | 120 | 16 |
| Drosopterin | 368.4 | 369.4 | 230.2 | 125 | 30 |
| Erythropterin | 265.1 | 266.1 | 220.1 | 110 | 8 |
| D-Neopterin | 253.2 | 254.2 | 206.2 | 115 | 14 |

**Table S7.**
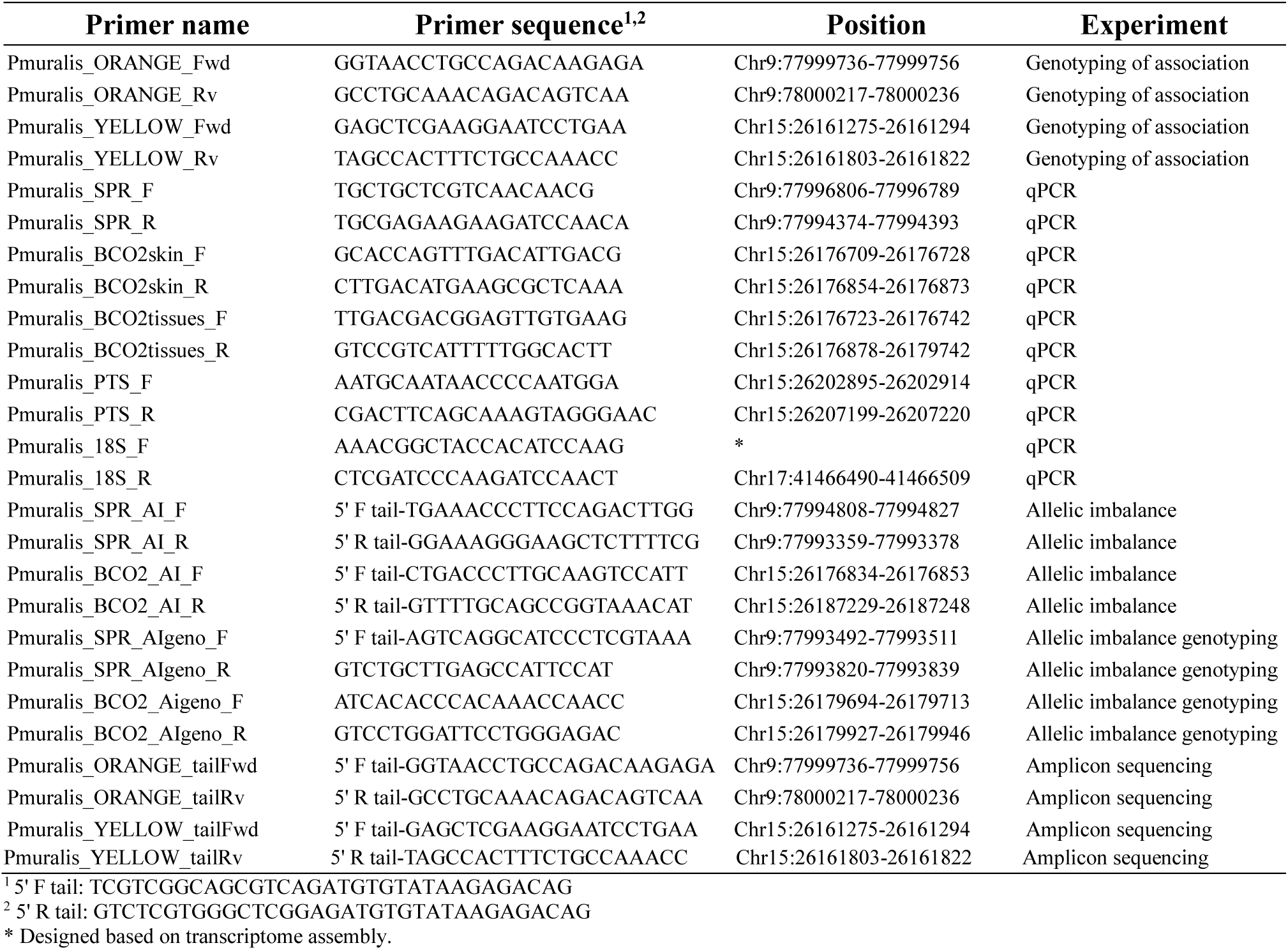
List of PCR primers that were used in this study for genotyping of the association, allelic imbalance, qPCR, and amplicon sequencing in *SPR* and *BCO2*

**Table S8.** Samples used for amplicon sequencing overlapping the regions of maximum association in *SPR* and *BCO2*

| **Sample code** | **Species** | **Morph** |
| --- | --- | --- |
| PY01 | *P. muralis* | Orange |
| TQ53 | *P. muralis* | Orange |
| PY03 | *P. muralis* | Orange |
| PC17 | *P. muralis* | Orange |
| PZ110 | *P. muralis* | Orange-white |
| TQ61 | *P. muralis* | Orange-white |
| TQ33 | *P. muralis* | Orange-yellow |
| VR01 | *P. muralis* | Orange-yellow |
| 2.5 | *P. muralis* | White |
| PZ115 | *P. muralis* | White |
| TQ45 | *P. muralis* | White |
| PY10 | *P. muralis* | White |
| 2.38 | *P. muralis* | Yellow |
| PZ13 | *P. muralis* | Yellow |
| TQ51 | *P. muralis* | Yellow |
| TQ57 | *P. muralis* | Yellow |
| 3.214 | *P. bocagei* | Orange |
| 3.216 | *P. bocagei* | Orange |
| 3.179 | *P. bocagei* | White |
| 3.19 | *P. bocagei* | White |
| 3.352 | *P. bocagei* | Yellow |
| 3.355 | *P. bocagei* | Yellow |
| DB26804 | *P. liolepis* | Orange |
| LLIVIA1 | *P. liolepis* | Orange |
| LLIVIA2 | *P. liolepis* | Orange |
| LLIVIA3 | *P. liolepis* | Orange |
| LLIVIA19 | *P. liolepis* | Orange |
| LLIVIA4 | *P. liolepis* | White |
| LLIVIA5 | *P. liolepis* | White |
| RIUCERD4 | *P. liolepis* | Yellow |
| DB20827 | *P. melisellensis* | Orange-white |
| DB20835 | *P. melisellensis* | Orange-white |
| DB20822 | *P. melisellensis* | Orange-yellow |
| DB20824 | *P. melisellensis* | Orange-yellow |
| DB20828 | *P. melisellensis* | White |
| DB20817 | *P. melisellensis* | Yellow |
| DB20818 | *P. melisellensis* | Yellow |
| DB26186 | *P. sicula* | Orange |
| DB26195 | *P. sicula* | Orange-white |
| DB26231 | *P. sicula* | Orange-white |
| DB26154 | *P. sicula* | White |
| DB26171 | *P. sicula* | White |
| DB26200 | *P. sicula* | White |
| 7.22 | *P. vaucheri* | Orange |
| 7.41 | *P. vaucheri* | Orange |
| 7.9 | *P. vaucheri* | Orange |
| 7.12 | *P. vaucheri* | White |
| 7.16 | *P. vaucheri* | White |
| 7.19 | *P. vaucheri* | White |
| 7.326 | *P. vaucheri* | White |
| 7.354 | *P. vaucheri* | Yellow |
| DB26532 | *P. wagleriana* | Orange-white |
| DB27409 | *P. wagleriana* | Orange-yellow |

**Table S9.**
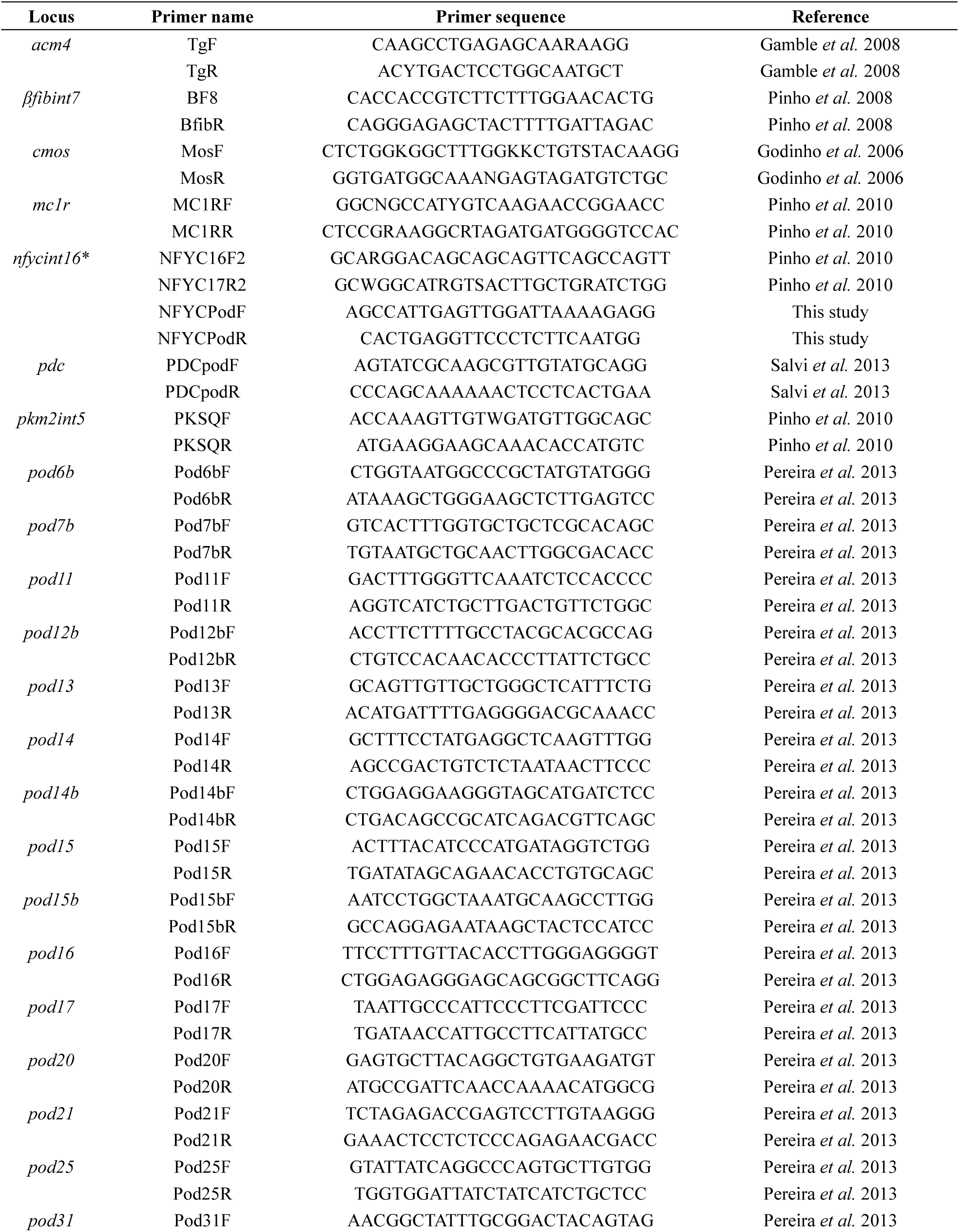
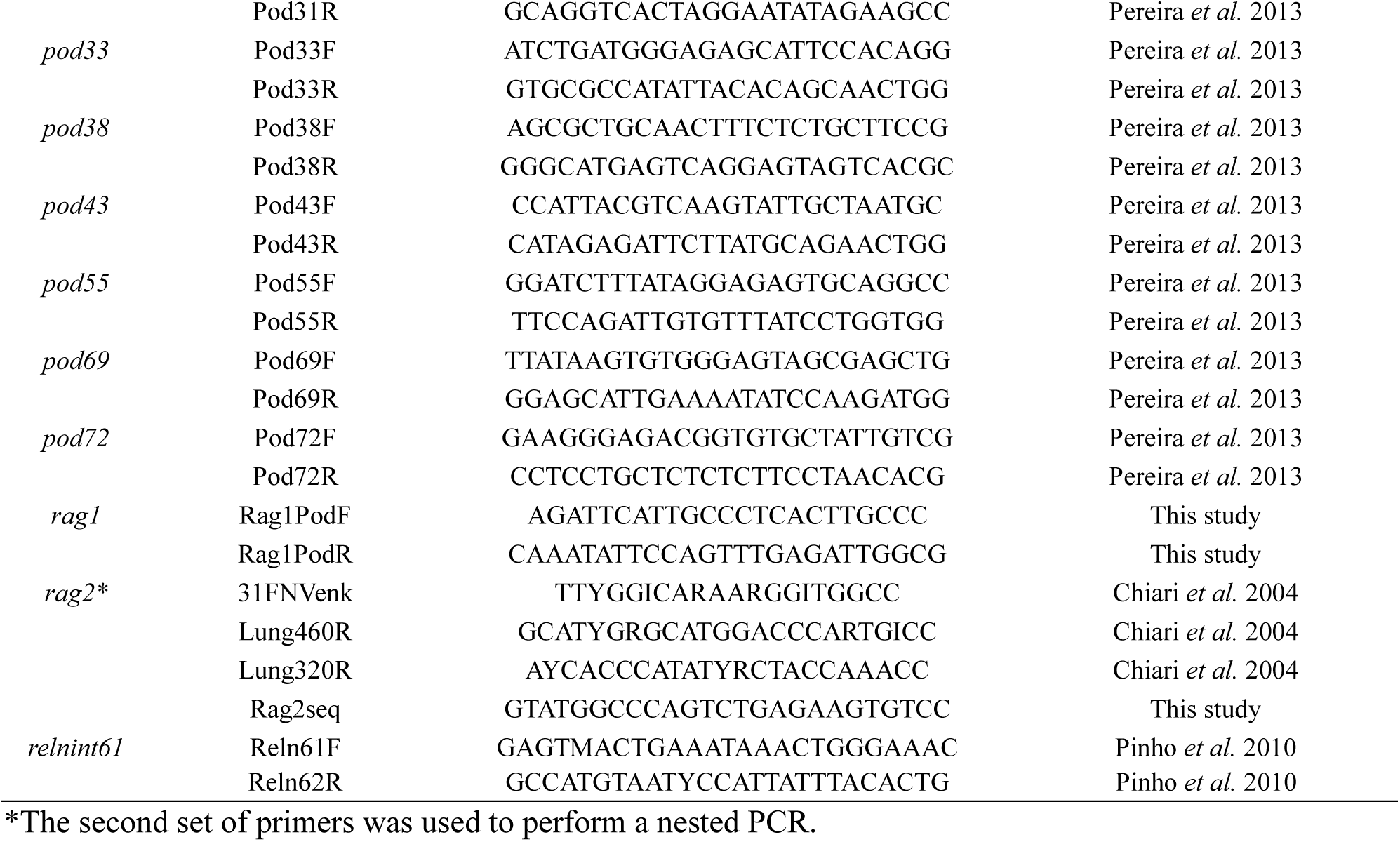
List of loci and primers used to amplify and sequence 31 loci randomly distributed across the genome in several color polymorphic species in the genus *Podarcis*

